# Synaptic dynamics are a tunable substrate sculpting neural population activity

**DOI:** 10.64898/2026.03.22.713064

**Authors:** Franziska Bender, B. Semihcan Sermet, Stefano Borda Bossana, Alessandro Barri, Selin Schamiloglu, Giovanni Diana, Maria-Miruna Costreie, Gael Moneron, Adam W. Hantman, David A. DiGregorio

## Abstract

A central tenet of cerebellar computation posits that granule cells generate sparse spatiotemporal activity patterns that support precisely timed motor and cognitive outputs. Using high-speed *in vivo* calcium and glutamate imaging combined with slice electrophysiology, we show that heterogeneous synaptic dynamics transform mossy fiber inputs into temporally sparse, sequential patterns of GC activity. Region-specific differences in MF glutamate release shape GC response duration and sparsity, tuning the temporal statistics of population sequences to match sensorimotor associative learning demands. These findings provide direct evidence for temporally sparse GC sequences and establish synaptic dynamics as a ubiquitous, tunable mechanism structuring neural activity in time.

## Introduction

Neural population activity across the brain unfolds as temporally structured sequences that underlie perception, memory, and precisely timed motor behaviors (*1–14*). A central organizing principle of these representations is sparsity: restricting activity to a fraction of neurons enhances storage capacity, robustness, and separability of neural codes (*2, 15–20*). In the cerebellum, the seminal theories of David Marr and James Albus proposed that expansion of mossy fiber (MF) inputs onto a vastly larger population of granule cells (GCs) generates sparse, high-dimensional representations for pattern separation (*16, 21–23*). This framework was later extended to the temporal domain, proposing that diverse GC response patterns form a high-dimensional temporal feature space that can be linearly decoded by Purkinje cells to produce precisely timed outputs (*24–27*).

Despite the decades of influence, direct experimental evidence of these theories has been lacking, in part due to limitations in the temporal resolution of typical calcium indicators (*28*) and the inability to resolve GC units with high-density electrophysiology (*29*). In particular, it remains unknown whether GC activity is temporally sparse and additionally contributes to pattern separation (*30, 31*). Work in other brain regions shows that temporally sparse population responses that tile time form an efficient substrate for downstream linear decoding (*7, 9*), and that sequence timescales are tuned to computational demands of different brain regions (*7*). However, whether such principles apply to the cerebellar GC layer remains unresolved.

Despite the ubiquity of neural sequences throughout the brain (*14*), a major outstanding question is therefore what circuit mechanisms generate temporally sparse neural sequences. Proposed explanations include structured connectivity, recurrent dynamics, and intrinsic cellular heterogeneity (*2, 7, 32–34*). Because short-term synaptic plasticity (STP) is ubiquitous across the nervous system (*35*) and easy to modulate (*36*), diversity in STP represents a compelling and general mechanism for shaping temporally structured population activity and its regional specialization. We therefore explored whether the known diversity of MF-GC synapses could be a mechanism to control the temporal statistics (sparsity and tiling) of GC population responses.

Here, we combine ex vivo brain slice recordings with high-speed *in vivo* two-photon imaging using ultrafast calcium and glutamate sensors to show that regional diversity in MF–GC STP provides a synaptic mechanism that converts dense afferent activity into temporally sparse GC sequences. By tuning synaptic timescales, this mechanism sets the temporal structure of GC activity to match the computational demands of distinct cerebellar regions.

## Results

### GC neural sequences form temporally sparse sensory representations

To determine whether cerebellar granule cells (GCs) form a temporal basis set resembling a neuronal activity sequence, we performed high-speed two-photon imaging of intracellular calcium responses evoked by air puff stimulation of the mouse whisker field (1-s air puff at 5 psi; **Fig. 1A, fig. S1A**). Resolving GC population dynamics at sub-second resolution *in vivo* is challenging, as GC spiking is inaccessible to modern silicon probes (*29*), and older genetically encoded calcium indicators blur rapid temporal structure (*37–39*). We therefore used the fast calcium indicator GCaMP8f (*40*) to resolve stimulus-locked GC population activity in Crus I. Whisker–air-puff–evoked responses were elicited using stimulus parameters similar to those used as conditioned stimuli in timing tasks (*41*), for which GC population dynamics have been hypothesized to provide a temporal basis for downstream learning (*27*). Whisker air puff produced calcium responses in ∼2% of identified GCs (**fig. S1B**), consistent with theoretical predictions that GCs form a sparse population representation of sensory-motor features (*16, 42, 43*).

**Fig. 1.**
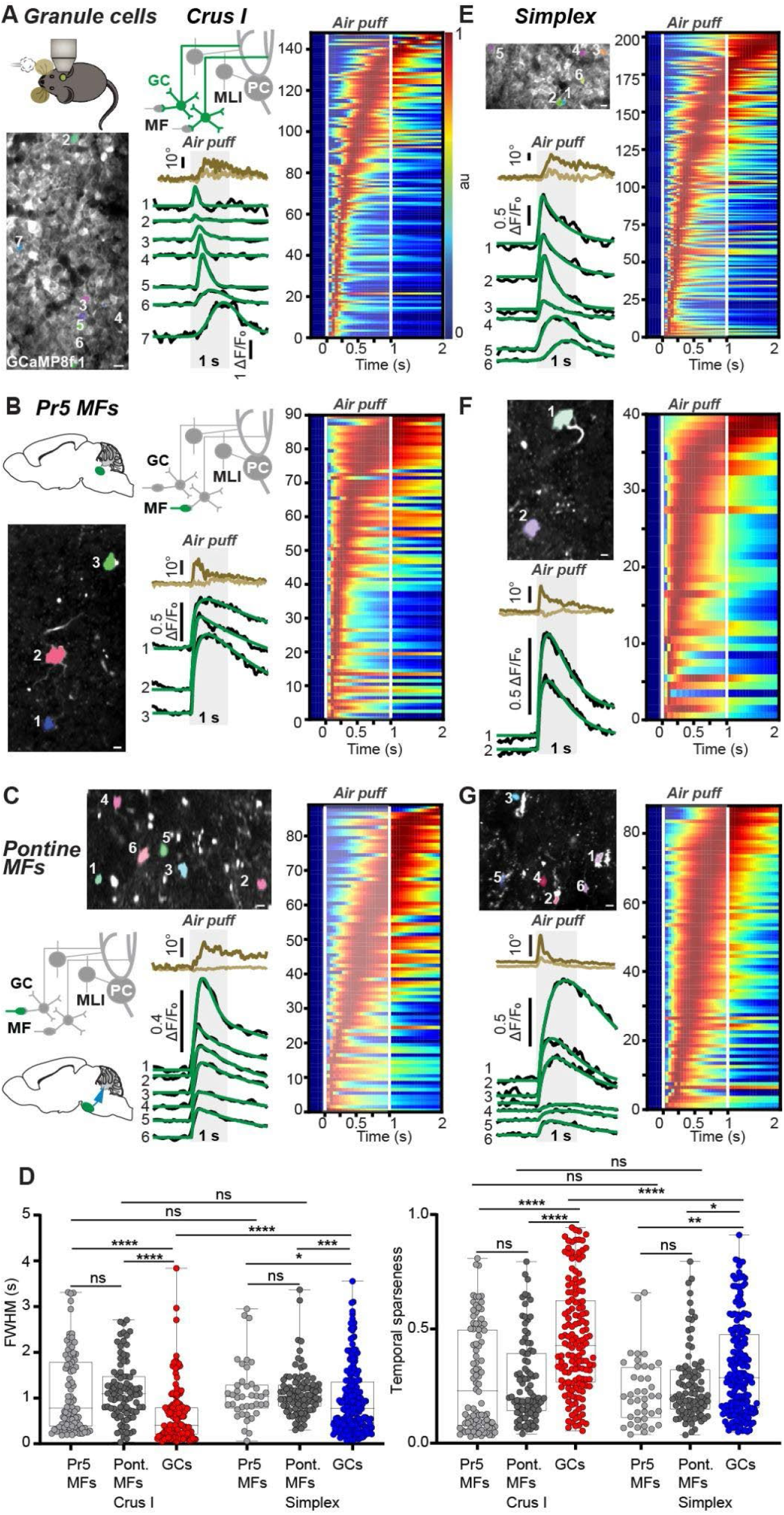
Crus I and Simplex mossy fiber and granule cell temporal representations of whisker air puffs. **(A)** Left, two-photon imaging of Crus I granule cells (GCs) expressing GCaMP8f. Center, whisker-air-puff responses of seven simultaneously recorded GCs averaged over 30 trials; the 1-s air puff is indicated by grey shading. Responses are shown above (light brown, ipsilateral; dark brown, contralateral to air puff). Right, average Crus I GC responses across experiments *(N* = 10 mice, *n* = 148 cells) shown as raster plots, normalized and sorted by time to peak. MLI: Molecular layer interneuron. PC: Purkinje cell. (**B and C**) Crus I mossy fiber (MF) responses to whisker air puffs from Pr5 (B; N = 4 mice, n = 90 cells) and pontine inputs (C; N = 3 mice, n = 89 cells). **(D)** Left, in Crus I and Simplex, response duration (full width at half maximum, FWHM) was greater in Pr5 (1.12 ± 0.09 s, *P* < 0.0001; Pr5 MFs, 1.13 ± 0.10 s, *P* = 0.027) and pontine MFs (1.16 ± 0.07 s, *P* < 0.0001; 1.12 ± 0.06 s, *P* = 0.0003) than in GCs (0.62 ± 0.05 s; GCs, 0.95 ± 0.05 s), with no difference between MF pathways *(P* = 0.17; *P* = 0.86). Crus I GC responses were shorter than Simplex GC responses *(P* < 0.0001), whereas Pr5 and pontine MF responses did not differ between regions *(P* = 0.80 and *P* = 0.34, respectively). Right, GC responses were temporally sparser than MF responses in Crus I (GCs, 0.46 ± 0.02; pontine MFs, 0.27 ± 0.02, *P* < 0.0001; Pr5 MFs, 0.28 ± 0.02, *P* < 0.0001) and Simplex (GCs, 0.33 ± 0.01; pontine MFs, 0.29 ± 0.02, *P* = 0.017; Pr5 MFs, 0.23 ± 0.02, *P* = 0.004). No difference was observed between Pr5 MFs and pontine MFs in Crus I *(P* = 0.1) or Simplex *(P* = 0.09). GC responses were more temporally sparse in Crus I than in Simplex *(P* < 0.0001), whereas no difference between regions was observed in MFs (Pr5, *P* = 0.86; pontine, *P* = 0.72). (**E–G**) Simplex responses to whisker air puffs: GCs (**E**; N = 8 mice, n = 203 cells), Pr5 MFs (**F**; N = 3 mice, n = 40 cells), and pontine MFs (**G**; N = 3 mice, n = 88 cells). Scale bars, 5 µm. Data are shown as median ± interquartile range, values in text as mean ± SEM. Statistical significance: *P* < 0.05 (*), *P* < 0.01 (**), *P* < 0.001 (***), *P* < 0.0001 (****).

We next examined the temporal characteristics of calcium responses within this sparsely recruited GC ensemble. GC responses were transient and generally shorter than the stimulus duration. They reliably produced peak response times that tiled the stimulus duration and were consistent across trials (**fig. S1**), reminiscent of a neural sequence. Individual GC responses, both within a single field of view and across animals, occurred as discrete calcium responses. Onset times spanned nearly an order of magnitude (10 ms–0.9 s). Response durations were heterogeneous (full-width-at-half-maximum (FWHM): 0.05–3.8 s; **Fig. 1**). At the population level, the temporal representation was biased toward GCs generating brief calcium transients near whisker air puff onset, whereas responses peaking later in the stimulus were progressively broader and less numerous. Consistent with this structure, peak response latency was positively correlated with burst duration (**fig. S2A**). We also observed a similar sequence-like population activity in GC axons within the molecular layer (**fig. S2B-D**). This confirmed that temporally sparse activity sequences found in the GC layer are also present across the population of GC axons likely to synapse onto a Purkinje cell (PC). In contrast, sequence-like population activity was not observed in molecular layer interneurons (MLIs), which exhibited shorter onset times and times-to-peak (**fig. S2E, F**). This suggests that sequence-like population activity is specific to GCs. Overall, results demonstrate that GC population activity is organized in a sequence-like manner at the sub-second time scale and suggest that diverse GC population activity can serve as a temporal basis set, as proposed by longstanding theories of cerebellar learning (*27*).

### GC neural sequence temporal sparsity requires a local transformation of MF input

The observed GC sequences could be inherited from their mossy fiber (MF) inputs or instead arise from a transformation of those inputs. To distinguish between these possibilities, we monitored MF activity projecting from the whisker somatosensory and motor cortices through the basal pons (secondary somatosensory inputs (*44*)) or directly via the trigeminal nucleus principalis (Pr5; primary somatosensory inputs, **Fig. 1B,C, fig. S3**) (*45–47*) to Crus I using the same calcium indicator and whisker stimulus. The fraction of activated MFs ranged from 16% in pontine MFs to 32% in Pr5 MFs (**fig. S1B**), clearly denser than the GC population (2%). Calcium responses were mostly excitatory, but we also observed inhibitory responses (Crus I GCs: 29% of responsive cells, Crus I pontine MFs: 5%, Crus I Pr5 MFs: 4%) and a small fraction of “off” responses, except in Pr5 MFs (Crus I GCs: 2%, Crus I pontine MFs: 1%, **fig. S4**). We focused subsequent analysis on excitatory responses.

The time course of MF responses was consistently slower than GC responses, exhibiting broader FWHM values (**Fig. 1D, left**) corresponding to a lower temporal sparsity, as quantified using a lifetime sparsity metric (**Fig. 1D, right**) (*48*). Cross-validation of MF and GC calcium responses to whisker air puffs demonstrated that responses in both populations were robust and reliable (**fig. S1**). The kinetic differences were not attributable to the calcium indicator, as *ex vivo* recordings showed that single- and burst-evoked calcium responses in MF terminals and GC somata had comparable rise and decay times (**fig. S5**). The FWHM of MF or GC calcium responses did not correlate with whisker air puff response duration (**fig. S6A-C**). Hence, GC population activity primarily reflected a sparse temporal code of the whisker air puff rather than motor-related responses. No correlation was observed between maximal deflection and response duration in GCs, Pr5 MFs, or pontine MFs (**fig. S6C**), further supporting a predominantly sensory representation. Analysis of the response onset variability showed that GCs exhibit similar heterogeneity (coefficient of variation, CV = 1.5) to that of their pontine MFs (CV = 1.6, two-sample Kolmogorov–Smirnov test, D = 0.119, *P* = 0.43). Pr5 MF response onsets, in contrast, were earlier and less diverse than GCs (CV = 1.0, two-sample Kolmogorov–Smirnov test, D = 0.43, *P* < 0.0001, **fig. S6D**). The short response onset of Pr5 MFs and the heterogeneity in pontine MF onset times (**fig. S6D**) are consistent with the direct arrival of the sensory information from the trigeminal nucleus and the delayed cortical information arising from cortico-pontine projections (*49, 50*), respectively. Together, these results indicate that the GC layer itself performs a critical transformation, converting dense MF temporal representations of sensory input into a temporally sparse sequence of GC activity that can serve as a basis set from which precise output activity dynamics can be learned (*7, 9*).

### Region-specific timescales of GC sequences

Cerebellar circuits support behaviors with distinct temporal requirements, and synaptic plasticity rules are tuned across cerebellar regions to match these timing demands (*51, 52*), raising the possibility that MF–GC transformations are region-specific and shape the temporal structure of local neural sequences. To determine whether MF–GC transformations generate distinct region-specific GC sequences, we measured Simplex GC and MF activity in response to whisker air puff (**Fig. 1E-G**). We took advantage of the fact that Crus I and Simplex regions both receive projections from Pr5 and the whisker somatosensory cortex (through the BPN, **fig. S3**) (*45*). No difference was observed between Crus I and Simplex in the relative number of activated GCs (3% in Simplex, similar to 2% in Crus I, **fig. S1B**). When comparing the temporal statistics of GC sequences, we observed no differences in the response width or lifetime sparsity of MFs between Crus I and Simplex (**Fig. 1D**). In contrast, GC responses from Crus I were significantly shorter and had higher lifetime sparsity than those from Simplex (**Fig. 1D**). Thus, despite receiving MF inputs with similar temporal statistics, MF-GC transformations are region-specific, generating GC activity with greater temporal sparsity in Crus I than in Simplex.

### Heterogeneity of Crus I and Simplex MF-GC synaptic dynamics in acute brain slices

To investigate what may underlie the difference in temporal sparsity between Crus I and Simplex, we next directly tested whether MF-GC synapses exhibit diverse and region-specific short-term plasticity (STP, **Fig. 1**). While cerebellar cortical models incorporating heterogeneous MF–GC STP predict that step changes in MF firing can be transformed into diverse GC temporal basis sets (*53*), evidence for regional diversity is lacking. We therefore recorded unitary MF–GC AMPA receptor–mediated excitatory postsynaptic currents (EPSCs) in whole-cell voltage-clamp in acute cerebellar slices using a blind minimum-stimulation approach (*54*). EPSCs from both Crus I and Simplex were highly heterogeneous in amplitude, coefficient of variation (CV), and paired-pulse ratio (PPR) across connections (**Fig. 2A, B, fig. S7**), as previously reported in the vestibulo-cerebellum (*31*). Our previous theoretical work showed that the time course of MF–GC synaptic dynamics during presynaptic trains directly determines the temporal statistics of GC firing responses, including their duration (*53*). We therefore quantified the timescales of STP at MF–GC synapses in Crus I and Simplex by estimating a weighted decay time constant (***τ***_*syn*_) associated with the adaptation of EPSC amplitudes during prolonged presynaptic firing (*53*) (**Fig. 2C, Materials and Methods**). Across three stimulation frequencies, ***τ***_syn_ was faster in Crus I than in Simplex (**Fig. 2D, E**). Thus, diversity in the STP timescale provides a candidate mechanism for the MF-GC transformation and the differences in GC temporal sparsity we observe between Crus I and Simplex.

**Fig. 2.**
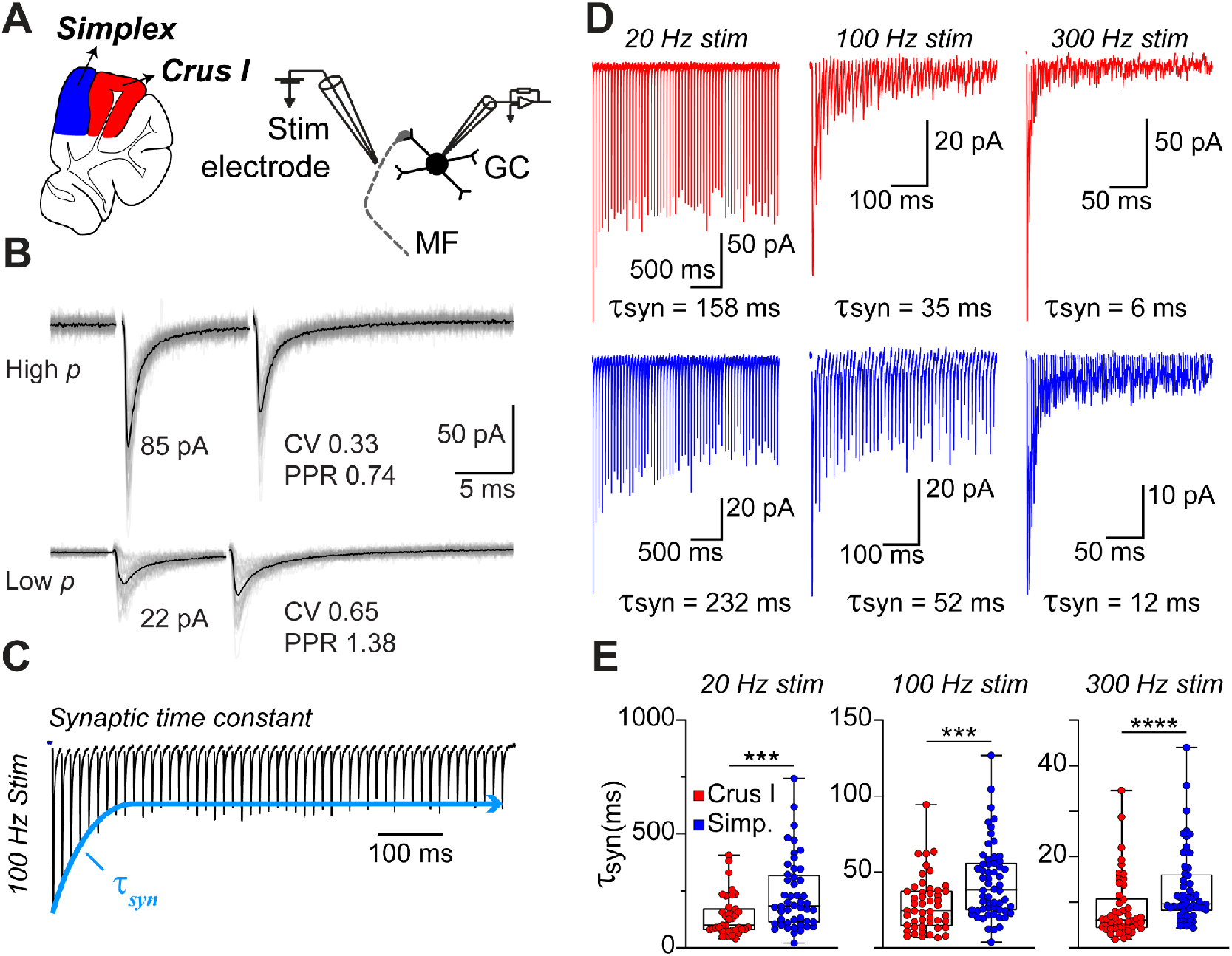
Regional differences in short-term synaptic dynamics at MF-GC synapses. **(A)** Schematic demonstrating the experimental design of whole-cell recordings from GCs in Crus I (red) or Simplex (blue) during MF stimulation in acute brain slices. **(B)** Unitary EPSCs from individual trials (gray traces) and mean (black trace) from two example MF-GC synapses with high (top) and low (bottom) release probability. EPSC peak amplitude, coefficient of variation of the peak amplitude (CV), and paired-pulse ratio (PPR) were measured for each synapse from responses to 100 Hz MF stimulation (see **fig. S7**). **(C)** Schematic illustrating exponential fit to EPSC peak amplitudes to extract synaptic time constants (***τ***_*syn*_). **(D)** Example EPSC traces (artifact subtracted and baseline corrected (see **Materials and Methods**) evoked by trains of MF stimulation at 20, 100, and 300 Hz in Crus I (top, red) and Simplex (bottom, blue). **(E)** Synaptic time constants (***τ***_*syn*_) at 20, 100, and 300 Hz in Crus I (red, *n* = 51) and Simplex (blue, *n* = 58). Data are presented as median with interquartile range. Crus I synapses display shorter ***τ***_*syn*_ than Simplex synapses at 20 Hz (Crus I: 98.85 ± 90.98 ms, Simplex: 183.58 ± 156.72 ms; *P* = 0.0007, Mann-Whitney test), 100 Hz (Crus I: 24.43 ± 18.04 ms, Simplex: 38.27 ± 24.54 ms; *P* = 0.0006, Mann-Whitney test), and 300 Hz (Crus I: 6.16 ± 6.81 ms, Simplex: 9.66 ± 8.10 ms; *P* < 0.0001, Mann-Whitney test) stimulation.

### MF–GC synaptic dynamics elicit diverse GC spiking patterns

Although regional differences in MF–GC synaptic dynamics provide a candidate mechanism for GC temporal diversity (*53*), other mechanisms could also contribute to the heterogeneity of whisker-air-puff-evoked GC responses, including Golgi-cell feedback (*27*) and tonic GABAergic inhibition, which influences the threshold for GC firing (*30, 55*). To isolate the contribution of synaptic dynamics alone, we first tested whether constant-frequency MF stimulation could elicit diverse GC spiking patterns in the absence of GABAergic inhibition. GC action potentials were recorded extracellularly during pharmacological block of both tonic and feedback GABA_A_ receptor–mediated inhibition (**Fig. 3A**). Despite identical presynaptic drive, individual GCs exhibited remarkably diverse firing dynamics (**Fig. 3B, C**). Most cells responded transiently to sustained 100 Hz MF stimulation, with firing rates that peaked rapidly and decayed with widely varying time constants (∼50–800 ms), closely matching predictions from our previous model (*53*) (**Fig. 3C**). Notably, a subset of GCs displayed delayed peak firing (>200 ms), demonstrating that synaptic dynamics alone can generate delayed spike responses, as previously observed for brief stimulus trains (*31*) (**Fig. 3B, magenta, orange**). Thus, intrinsic GC layer mechanisms can transform constant MF input into temporally structured spike output.

**Fig. 3.**
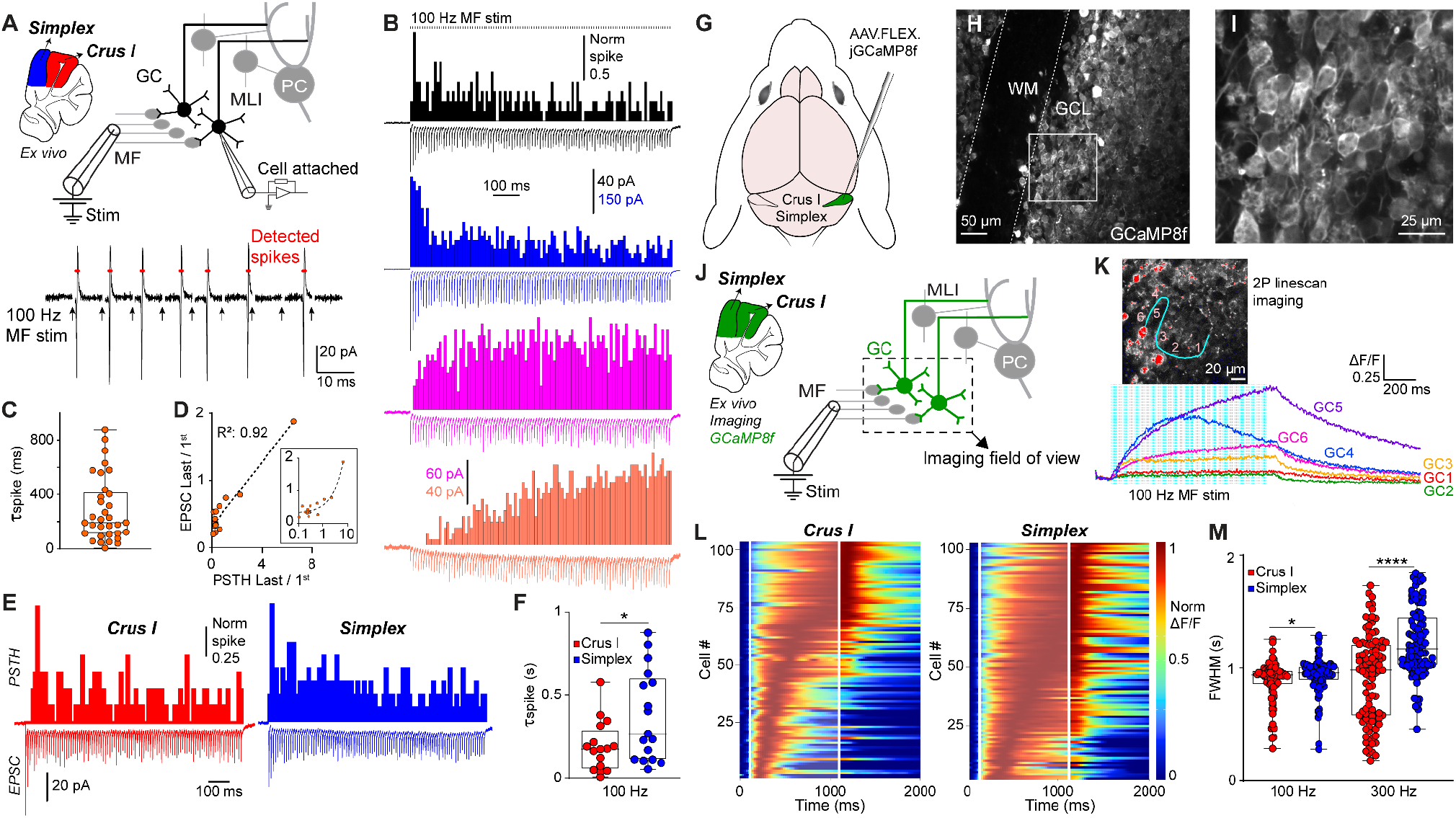
Synaptic properties shape GC temporal dynamics. **(A)** Top: Schematic of cell-attached GC recording during MF stimulation. Bottom: Example of 100 Hz MF stimulation and detected GC spikes. Arrows indicate stimulation times; stimulation artefacts were blanked. **(B)** PSTH (above) and mean EPSC (below) responses of four GCs following a 1 s, 100 Hz MF stimulus. Top, black vertical ticks indicate stimulation time points. **(C)** Quantification of the spike decay time constant (***τ***_*spike*_) of all cells responding to 100 Hz MF stimulation (204.81 ± 240.79 ms, *n* = 33 cells). **(D)** Correlation plot of synaptic EPSC ratio vs spike PSTH ratio *(n* = 16), measured as the ratio of the steady state EPSC amplitudes or PSTH counts (last) over the first EPSC response amplitude or PSTH bin count. The inset shows the horizontal axis on a log scale (regression line: *R*^*2*^ = 0.92). **(E)** Representative GC firing PSTHs (top) and average EPSCs (bottom) in Crus I (red, left) and Simplex (blue, right) during 100 Hz MF stimulation. **(F)** Quantification of GC spike decay time constant (***τ***_*spike*_) in Crus I (red, 178.06 ± 156.72 ms, *n* = 16) and Simplex (blue, 269.14 ± 272.51 ms, *n* = 17; *P* = 0.049, Mann-Whitney test). **(G)** Schematic of the AAV injection protocol to express GCaMP8f in Crus I and Simplex GCs. **(H)** Expression of GCaMP8f in GCs in Crus I. WM, white matter; GCL, granule cell layer. **(I)** Expanded version of the white-framed field of view in **H. (J)** Schematic of two-photon calcium imaging of GCs while stimulating MFs with a large-diameter glass electrode. **(K)** Example calcium responses of six GCs recorded using two-photon linescan imaging. The inset shows the imaging field of view and the scanning line drawn over 6 GCs. **(L)** Raster plots of Crus I (left) and Simplex (right) GC responses to 300 Hz MF stimulation. Each row represents the responses of a single GC. White lines indicate stimulus onset and offset. **(M)** Quantification of FWHM of GC responses to 100 and 300 Hz MF stimulation in Crus I (red, 0.93 ± 0.19 s, *n* = 80 for 100 Hz; 0.98 ± 0.38 s, *n* = 104 for 300 Hz) and Simplex (blue, 0.96 ± 0.18 s, *n* = 89 for 100 Hz; 1.17 ± 0.30 s, *n* = 103 for 300 Hz). Data are presented as median with interquartile range. FWHM is significantly shorter in Crus I at both 100 Hz (Mann-Whitney test, *P* = 0.03) and 300 Hz (Mann-Whitney test, *P* < 0.0001) MF stimulation.

We next asked whether GC firing diversity directly reflects heterogeneity in MF–GC synaptic dynamics. We reasoned that if synaptic inputs govern spike timing, the time course of EPSC dynamics during high-frequency stimulation (***τ***_*syn*_) should predict the temporal profile of GC spiking (***τ***_*spike*_). To test this, we recorded spike PSTHs from individual GCs in cell-attached mode before transitioning to whole-cell configuration to measure MF–GC EPSCs evoked by identical stimulus trains (**Fig. 3B**). To include GCs with slowly varying firing rates that deviate from exponential decay, we calculated EPSC and PSTH ratios as proxies for ***τ***_*syn*_ and ***τ***_*spike*_. Across cells, we observed a strong correlation between the two *(R*^2^=0.92), establishing a tight synapse-to-spike correspondence (**Fig. 3D**). Consistent with regional differences in MF-GC STP, GC spiking dynamics were significantly slower in Simplex than in Crus I (**Fig. 3E, F)**, demonstrating that region-specific synaptic dynamics set the temporal profile of GC output. These results identify STP heterogeneity as a primary determinant of GC firing diversity.

To determine if these single-cell mechanisms scale to the population level, we performed high-speed two-photon imaging of GCaMP8f-expressing GCs in acute slices during constant MF stimulation (**Fig. 3G-I**). Identical MF input elicited sequence-like GC population responses with a broad distribution of response latencies and durations (**Fig. 3K, L**). At stimulation frequencies comparable to whisker-air-puff-evoked MF firing in Pr5 (*56*) (100 and 300 Hz), GC responses in Crus I were significantly briefer than those in Simplex (**Fig. 3L, M**), recapitulating the regional differences observed *in vivo* (**Fig. 1**). These population-level dynamics mirror regional specializations in MF–GC STP (**Fig. 2**), supporting the idea that synaptic diversity directly organizes GC temporal sequences. Notably, blocking GABAergic transmission had minimal effect on these temporal patterns (**fig. S8**). Together, these single-cell and population experiments demonstrate that heterogeneity in MF–GC synaptic dynamics is sufficient to generate region-specific, temporally ordered GC firing patterns under constant input, even in the absence of feed-forward and feed-back inhibition.

### Network simulations with dynamic MF-GC synapses reproduce observed GC sparsification and region specificity

To test whether region-specific MF–GC synaptic dynamics are a plausible mechanism for generating sparse, region-specific GC representations, we leveraged a cerebellar cortex firing-rate model that incorporates experimentally constrained MF–GC STP diversity (*53*) (**Fig. 4A, B**). With minimal parameter tuning, regional differences were captured by selectively increasing the refilling time constant (***τ***_*ref*_) of the low-release-probability vesicle pool (**Fig. 4B**), resulting in Simplex-like synapses having a higher PPR and higher steady-state amplitudes than Crus I-like synapses, without affecting the mean initial EPSC, in qualitative agreement with our experimental findings (**Fig. 2** and **fig. S7C**). This also reproduced experimentally observed differences in synaptic time constants, yielding a 1.25–1.5-fold slowing of ***τ***_*syn*_ in Simplex (**Fig. 4 C, D**, see **Fig. 2G**). The model was driven with MF activity patterns approximating pontine and Pr5 whisker-air-puff-evoked inputs (**Fig. 1**, see **Materials and Methods**). Simulated transmitter release and GC population activity were markedly sharpened relative to MF input, as quantified by reduced response FWHM (**Fig. 4E, G**). Simplex-like synapses produced broader GC activity profiles than Crus I-like synapses yet remained substantially sharper than their parent MF inputs (**Fig. 4E, G**). In contrast, removing STP resulted in only modest sharpening (**Fig. 4F, G**), suggesting that MF– GC STP is the principal mechanism underlying region-specific and temporally sparse GC activity.

**Fig. 4.**
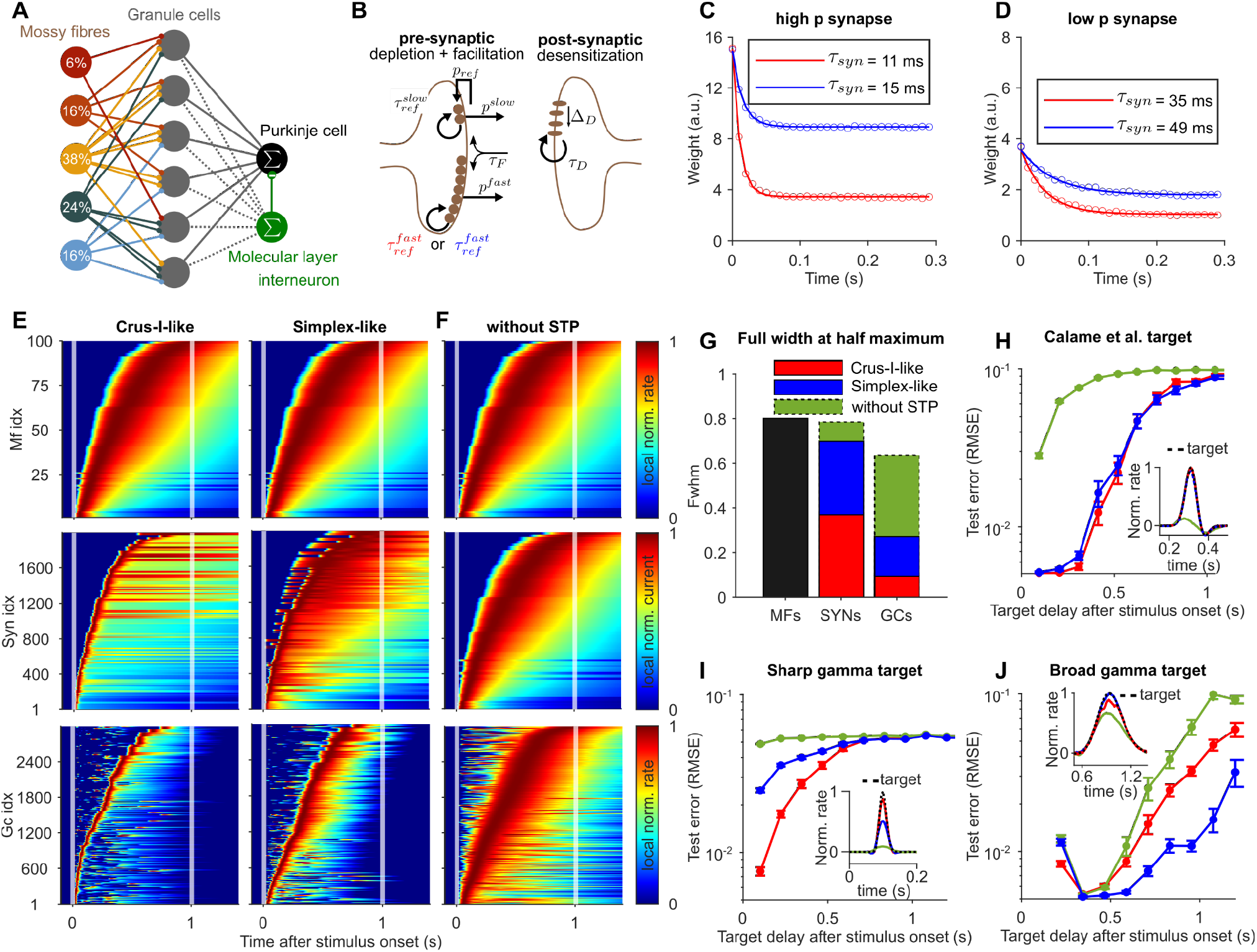
Model simulations show that region-specific heterogeneous synaptic dynamics can sparsify GC representations differently and introduce learning biases. **(A)** Scheme of rate-based cerebellar cortex model including MFs, GCs, one Purkinje cell (PC), and one molecular layer interneuron (MLI), adapted from (*53*). Colors and percentages indicate the relative frequency of MF-GC synapse types; see (*53*). **(B)** Scheme of MF-GC synaptic model showing the principal model parameters, adapted from (*53*). The time constant of the fast-refilling pool is changed to mimic Crus I (red) or Simplex (blue) synapses. **(C)** Simulated average synaptic EPSCs of a high-release-probability synapse in response to a 100 Hz stimulation for Crus I-like (red dots) and Simplex-like synapses (blue dots); solid lines: mono-exponential fits with respective time constants shown in the box. **(D)** Same as **C**, but for a low-release-probability synapse. **(E)** Simulated stimulus-evoked population activity of MFs (top), MF-GC synapses (middle), and GCs (bottom); color-coded MF and GC firing rates as well as MF-GC synaptic currents are individually normalized and sorted according to peak position. Left column: activities for Crus I-like synapses; right column: activities for Simplex-like synapses. **(F)** Same as **E** but for static MF-GC synapses and with an identical GC threshold as in **E. (G)** Average FWHM values for populations shown in **E** and **F. (H)** Reconstruction error of a brief temporal signal representing forelimb reach kinematics (*57*) at varying target delays for different GC basis sets; each point represents an average over 30 learning simulations. Inset: example of target signal (dashed black line) and simulated PC responses after learning. **(I)** Same as **H**, but for narrow gamma target signals with a SD of 0.01 s. **(J)** Same as **H**, but for broad gamma target signals with an SD of 0.15 s.

We next asked whether these region-specific GC temporal bases impose distinct constraints on downstream learning. A linear readout unit representing a Purkinje cell (PC; **Fig. 4A**) was trained, using a biologically plausible supervised learning rule, to generate predefined PC output activity profiles from simulated GC activity (see **Materials and Methods**). Incorporating dynamic MF–GC synapses substantially improved learning, enabling both Crus I- and Simplex-like models to reproduce realistic PC target responses (*57*) for target profile onset delays up to ∼700 ms (**Fig. 4H**). Strikingly, the two models exhibited complementary learning specializations. Crus I-like synapses preferentially supported learning of brief, early-onset PC targets (**Fig. 4I; fig. S9A-D**), whereas Simplex-like synapses were more effective for broader, late-onset targets (**Fig. 4J**; **fig. S9A-D**). These biases persisted under full gradient descent (**fig. S9A-C**), indicating that they arise from intrinsic differences in GC temporal basis structure rather than properties of the learning rule. In addition, when reducing synaptic diversity, learning performance in broad, late-onset targets was decreased in both Crus I- and Simplex-like models (**fig. S9E-H**). Together, these simulations indicate that MF–GC pre-synaptic dynamics can be regionally tuned to sharpen GC temporal representations and optimize learning to match distinct behavioral time scales.

### Region-specific GC dynamics are inherited from differences in regional neurotransmitter release dynamics

If MF–GC STP diversity underlies region-specific GC sequence sparsification, it should be directly reflected in the temporal dynamics of vesicular glutamate release, as shown by our simulations, in which vesicle depletion transforms sustained MF firing into brief, transient synaptic drive. Using a fluorescence-based genetically encoded glutamate sensor (*58*), we directly tested these predictions by monitoring MF glutamate release onto single GC dendritic claws. Expression of the sensor postsynaptically also allowed us to examine the within-cell diversity of neurotransmitter release at all incoming synapses. Because GCs are extremely densely packed, sparse expression is essential to resolve individual MF–GC synaptic connections. We therefore took advantage of a sparse tTA-conditional transgenic line (TCGO-dark), generated by CRISPR-mediated removal of the original YFP tag from the TCGO line (*59*), to express iGluSnFR-A184V in a small subset of GCs in Crus I and Simplex (**Fig. 5A,B**). The decay time of iGluSnFR responses to a single action potential is similar to that of GCaMP8f (20–30 ms (*58*)), indicating that differences in response dynamics cannot be attributed to sensor kinetics. In contrast to GCaMP8f responses, which were a proxy for MF activity, changes in iGluSnFR fluorescence in GC dendritic claws during whisker air puff stimuli reflect short-term alterations in spike-evoked glutamate release from individual MF terminals onto GC dendrites (**Supplementary Movie S1**).

**Fig. 5.**
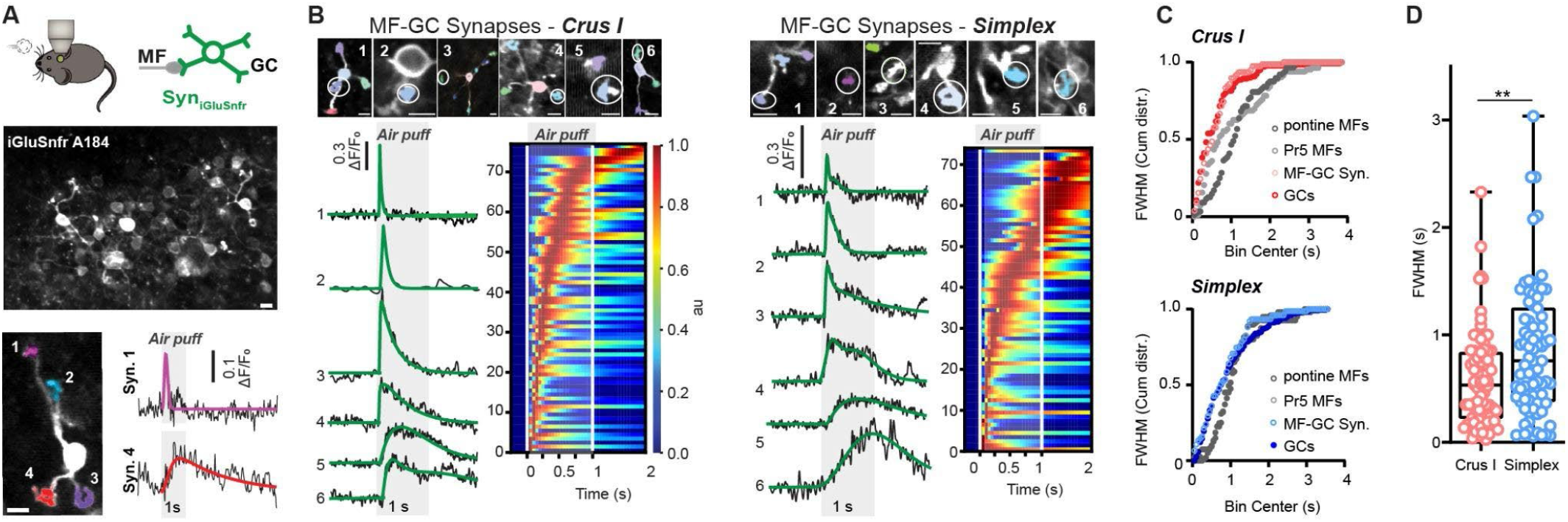
*In vivo* MF neurotransmitter release dynamics are diverse and faster than MF activity, providing a substrate for precise temporal learning. **(A)** Schematic of experimental setup (top). Crus I GCs labeled sparsely with GluSnFR under the TRE promoter in a TCGO mouse (middle). Note that single synapses can easily be identified. Representative example of two MF synapses onto the same GC with different responses (bottom). **(B)** Representative examples of Crus I (left) and Simplex (right) whisker air puff responses from different MF-GC synapses and cells. Note variability across responses. Raster plots of responses of all measured synapses averaged across trials and normalized on the right, respectively (Crus I: *N* = 9, *n* = 78, Simplex: *N* = 8, *n* = 75). **(C)** Cumulative distribution of FWHM for Pr5 MFs, pontine MFs, GCs, and MF-GC synapses. Note the overlap of GC and MF-GC synapse responses, indicating that GCs mimic the temporal response distribution of synaptic responses (two-sample Kolmogorov–Smirnov test, Crus I D = 0.10, *P* = 0.66, Simplex D = 0.13, *P* = 0.27), while it differed between MF-GC synapses and pontine MFs (Crus I D = 0.47, *P* < 0.0001, Simplex D = 0.29, *P* = 0.0014) or Pr5 MFs (Crus I D = 0.86, *P* = 0.028, Simplex D = 0.30, *P* = 0.017). **(D)** The difference in GC whisker air-puff response FWHM between Crus I mimics the difference in MF-GC synapse GluSnFR release between Crus I (0.58 ± 0.05 s) and Simplex (0.85 ± 0.07 s; *P =* 0.0049). Scale bars, 5 µm. Data are shown as median ± interquartile range; values in the text are mean ± SEM.

Consistent with heterogeneous presynaptic release dynamics, iGluSnFR responses recorded from different claws of the same GC could differ markedly (**Fig. 5A**), mirroring the diversity of synaptic strength and STP observed at MF–GC synapses in brain slice experiments (**Fig. 3**) (*31*). Glutamate responses to repeated presentations of the same whisker stimulus were robust across trials in both Crus I and Simplex (**fig. S10A**). Like GC activity responses, neurotransmitter release response amplitudes were not correlated with whisker air puff responses (**fig. S10**). The time course of the glutamate release was diverse in both Crus I and Simplex and reminiscent of neural sequences in GCs (**Fig. 5B**). The distribution of FWHM of MF-GC synaptic responses was more similar to that of GCs for both Crus I and Simplex than to that of pontine MFs or Pr5 MFs (**Fig. 5C**). Similar to GC neural sequences (**Fig. 1**), glutamate release at MF-GC synapses was significantly briefer in Crus I than in Simplex (**Fig. 5D**), consistent with the finding that synapses in Crus I were faster than those in Simplex. Moreover, glutamate release in Simplex was less temporally sparse than in Crus I, similar to GCs (**fig. S10**). Together, these results demonstrate that sparse, region-specific GC sequences arise from temporally structured neurotransmitter release at MF–GC synapses driven by presynaptic STP diversity.

## Discussion

Neural sequences are a ubiquitous coding strategy throughout the brain, yet the circuit and synaptic mechanisms that generate them remain poorly understood. Here, we identify a synaptic mechanism by which the cerebellar input layer transforms sustained sensory drive into temporally sparse population activity that forms a neural sequence tiling the stimulus duration. Cerebellar granule cells (GCs) have long been hypothesized to generate diverse temporal representations of contextual information, which are essential for associative motor learning (*24, 27*). Using high-speed multiphoton calcium imaging, we showed that air puff stimulation of the whisker field evoked transient GC responses whose peak amplitudes tiled the stimulus, producing a GC activity sequence with narrower temporal profiles than the calcium responses of their mossy fiber (MF) inputs, and that this sharpening is region-specific. Combining *ex vivo* electrophysiology, network modeling, and *in vivo* glutamate imaging, we demonstrated that heterogeneous neurotransmitter release dynamics at MF–GC synapses drive the temporal sharpening of GC activity, are region-specific, and enhance the precision of learning specific time-dependent output activity dynamics. Together, these findings provide direct evidence that heterogeneous synaptic dynamics are a tunable substrate that can sculpt population activity over time to meet the precise timescale demands of cerebellar region-dependent behaviors.

GC neural sequences exhibit multiple computationally relevant properties. Efficient learning systems benefit from internal representations in which complex signals are encoded as combinations of sparsely active units, greatly expanding representational capacity in neural circuits and artificial networks (*60, 61*). In the cerebellum, sparse GC representations are essential for efficient pattern separation (*18, 21, 22*). Our results show that such spatial sparsity is achieved through expansion recoding (*30, 42, 43*), whereby fewer than 2% of identified GCs respond to a simple sensory stimulus, compared with a maximum of 16–32% of identified MFs. This spatial sparsity is accompanied by pronounced temporal sparsity within the responsive GC ensemble, providing an additional mechanism for discriminating dynamic patterns of GC activity, as we previously demonstrated for GC delay coding (*31*). The minimal temporal overlap of GC responses yields a population sequence that approximates an orthogonal representation of time, facilitating the learning of time-dependent outputs (see **Fig. 4** and (*7, 9, 62*)). The latency of peak GC activity scaled linearly with response width (**Fig. 1**; **fig. S2**), consistent with Weber’s scalar timing principle, where variability time estimates increase linearly with the time interval (*63, 64*). Such Weber-like scaling is a key requirement of theoretical models proposing that cerebellar circuits perform Bayesian estimation of time intervals by learning prior distributions of temporal intervals (*53, 62*). Our results, therefore, provide experimental evidence that GC temporal representations satisfy a central prediction of Bayesian models of cerebellar timing. Together, these findings underscore how high–temporal-resolution measurements of neural activity within defined circuit layers can reveal circuit-level computational principles.

Heterogeneous STP sculpts the temporal sparsity of GC neural sequences, thereby expanding the capacity of cerebellar circuits to encode sensory dynamics and support precisely timed motor outputs. Because vesicle replenishment kinetics influence the time course of STP and are subject to neuromodulatory control (*65, 66*), the dynamic regulation of STP provides a plausible route for adapting the temporal properties of GC activity (*36*) to the behavioral context without altering circuit connectivity. Region-specific synaptic dynamics provide a substrate by which different cerebellar regions could adjust the precision and timescales of temporal learning to be in line with known regional differences in learning rules (*51*). The mechanisms that establish region-specific synaptic timescales could arise through genetic specification, developmental learning, or activity-dependent homeostatic regulation. Regardless of their origin, these differences bias learning in predictable ways: brief GC responses preferentially support learning of fast PC target signals (**Fig. 4H, I**), whereas broader GC responses better support slower target dynamics (**Fig. 4J**). Such biases may align with a mediolateral gradient in cerebellar function, spanning rapid sensorimotor processing to more integrative cognitive operations (*67*). Alternatively, the whisker sensorimotor representations in Crus I (*47, 68, 69*) may require fast, temporally sparse representations to support rapid deflections and high spatial acuity, whereas the Simplex region, involved in limb motor control (*70*), may benefit from slower temporal representations to drive motor learning. Notably, the longer GC responses observed in Simplex may also promote generalization by enabling broader, more overlapping temporal representations, potentially accelerating learning across related tasks. Thus, the regional diversity and modulation of MF-GC synaptic dynamics may provide a flexible strategy for the cerebellum to specifically balance temporal precision, bandwidth, and generalization for learning specific behaviors.

Heterogeneity in short-term synaptic plasticity is conserved across animal phyla, from invertebrates to humans (*35*), and implicated in shaping neuronal dynamics and computations (*71–74*). Our results position STP diversity within a broader population-level framework by showing that heterogeneous synaptic dynamics provide a biophysical substrate for transforming sensory input into temporally sparse neural sequences tuned for task-specific computations. Given the ubiquitous nature of STP across neural systems, synaptic heterogeneity may be a fundamental circuit feature that shapes the timing, duration, and distribution of population activity. Such diversity could therefore be tuned to optimize associative learning beyond the cerebellum and influence a wide range of time-dependent computations across sensory, motor, and cognitive domains.

## Funding

Work in the DD laboratory was supported by the Fondation pour la Recherche Médicale (FRM EQU202003010555), Fondation pour l’Audition (FPA-RD-2018-8), the Agence Nationale de la Recherche (ANR-17-CE16-0019, ANR-18-CE16-0018, ANR-19-CE16-0019-02, ANR-21-CE16-0036-01, ANR-23-CE37-0026-03, and ANR-23-CE23-0034-03), and by the NIH (R21 EY026434). FB was additionally supported by the Human Frontier Science Program postdoctoral fellowship (HFSP, LT000674/2019-L), BSS by Pasteur-Roux-Cantarini postdoctoral fellowship from the Institut Pasteur (PRC), SS by the NIH (F32 NS141896), and SBB by a stipend from the Pasteur-Paris University (PPU) International PhD Program. We thank Maureen McFadden for helping with *ex vivo* synaptic time constant analysis. We thank Florian Ruckerl for technical assistance. We thank the Wang lab for providing the AAVDJ-CAG-FLEx-jGCaMP8f construct. We thank the Histology Platform at the Institut Pasteur for providing high-resolution histology images.

## Contributions

D.A.D., A.H., F.B., and B.S.S. designed experiments. F.B. performed and analyzed *in vivo* experiments. B.S.S. performed and analyzed *ex vivo* experiments. S.B.B. performed and analyzed *ex vivo* voltage-clamp and part of the cell-attached experiments. A.B. designed and performed model simulations. S.S. performed part of *the in vivo* GC Calcium imaging experiments. M.M.C. performed MLI *in vivo* experiments. G.D. and G.M. conceived the behavioral analysis pipeline. A.H. provided viral constructs for *in vivo* synapse imaging. All authors wrote and reviewed the manuscript.

## Competing interests

The authors declare no competing interests.

## Materials and Methods

### Mice

All animal experimental procedures complied with the ARRIVE guidelines and were conducted in compliance with French and European regulations on care and protection of laboratory animals (EC Directive 2010/63, French Law 2013-118, February 6th, 2013) and approved by the Ethics Committee for animal experimentation at the Pasteur Institute (CETEA), or the Institute of Experimental Medicine Protection of Research Subjects Committee, or the University of Colorado Anschutz Medical Campus Institutional Animal Care and Use Committee. Mice were housed in the Pasteur Institute animal facilities, accredited by the French Ministry of Agriculture for performing experiments on live rodents, or at the University of Colorado Anschutz Medical Campus. Mice were kept on a reverse 12 h or 14:10 h light/dark cycle with food and water *ad libitum*. For *in vivo* experiments, GC data came from 14 Gabra6^tm2(cre)Wwis^ (Jackson Laboratory strain 3047798) and 8 B6.D2-Tg(Gabra6-cre)B1Lfr/Mmucd mice, MF data from 9 αGabra6-Cre (B6J-Duke) mice, GC-MF Synapse data from 29 TCGO dark mice (*59*), and MLI data from 7 WT (C57BL/6J, Jackson Laboratory strain 000664) mice. For *in vitro* calcium imaging experiments, data came from 14 alphaGabra6(cre)(B6j-Duke) mice. For *in vitro* synaptic recordings, data were obtained from F1 mice (N = 103, CB6F1 cross of BALB/cJ and C57BL/6J).

### Viral injections

For viral injections, mice were first deeply anesthetized with 1 to 4% isoflurane mixed in oxygen. They were then placed in a stereotaxic surgery frame using a mouth clamp. Carprofen (Metacam) was injected intraperitoneally as an analgesic, and mice received additional subcutaneous injections of buprenorphine and dexamethasone. After repeatedly disinfecting the top of the mouse head with liquid betadine and 70% ethanol, lidocaine was injected under the scalp for local anesthesia. A heating blanket with a closed-loop temperature control system was used to maintain body temperature at ∼37 °C. Throughout the surgery, temperature and breathing were monitored closely. Eyes were covered with ointment (Viscotears, Alcon, USA; VITA-POS, Pharma Medica AG, Switzerland) to prevent drying. Small scissors were used to open the scalp and expose the skull. Cotton swabs and a scalpel were then used to clean the skull and remove remaining tissue. Lateral muscles on the skull were detached using a scalpel to access the cerebellum. After careful alignment of the skull on the stereotaxic frame, we identified the region of interest for injections. For all targeted structures, coordinates were measured from bregma as follows: Pr5, 5.0 mm posterior, 1.8 mm lateral, 3.5 mm deep from brain surface; BPN, 4.0 mm posterior, 0.4 mm lateral, 5.8, 5.5, 5.2, and 5.0 mm deep from brain surface; Crus I, 6 mm posterior, 3 mm lateral, 0.3-0.7 mm deep from brain surface; Simplex, 6 mm posterior, 1.75 mm lateral, 0.3-0.7 mm deep from brain surface. A small craniotomy of about 0.5 mm diameter was made at the targeted location, and forceps were used to lift the bone cap to access the brain. A thin glass pipette (PCR Micropipets 1-10 µl, Drummond Scientific Company, USA) was first pulled, and then the tip was broken using a tissue to give a 21–27 µm inner tip diameter. The pipette was filled with mineral oil, and then the tip was filled with the AAV vector. The pipette was slowly lowered to the target location in the brain, and injections were performed using an oil-hydraulic injector (Nanoject III, Harvard Apparatus). To specifically express GCaMP8f in GCs, we used the Gabra6-Cre mouse line, which expresses Cre recombinase exclusively in GCs (*75*). We delivered AAVDJ-CAG-FLEx-jGCaMP8f (virus titer: 5×10^13^ GC/ml, 1:10 diluted in NaCl), a gift from Wang laboratory (Princeton University), to cerebellar Crus I and/or Simplex at a speed of 2 nL/s, 50 nL per injection site to image GC somas and PFs. In a subset of experiments, we targeted GCs by delivering AAV9-syn-FLEX-jGCaMP8f-WPRE (Addgene, virus titer: 3×10^13^ GC/ml, undiluted) in B6.D2-Tg(Gabra6-cre)B1Lfr/Mmucd mice to Crus I and/or Simplex (0.3 and 0.6 mm deep from brain surface), at a speed of 1 nL/s, 150 nL per injection site. To express GCaMP8f in Pr5 and BPN, we used AAV1-hSyn-gCaMP8f (3×10^13^, 1:5 diluted in NaCl). The virus was delivered at 2 nL/s, 50 nL per injection site. For MLI experiments, AAV1-syn-GCaMP8f-WPRE (virus titer: 3 x 10^13^ GC/ml, 1:10 diluted in NaCl, from the Adam Hantman laboratory, Janelia Research Campus) was delivered at 2 nL/s, 100 nL at 0.2 mm depth. For the sparse expression of SF-iGluSnFR in GCs, we injected AAVDJ.TRE.SF-iGluSnFR.A184V (virus titer: 9.32×10^12^ GC/ml, 1:10 diluted in NaCl (*58, 76*)) in TCGO (*59*) mice that express tetracycline transactivator in a subset of GCs. Injected mice were then kept for 2 weeks to allow transgene expression.

### Headplate and cranial window surgery for *in vivo* imaging experiments

Animals were given buprenorphine (Buprecare, 0.05-0.1 mg/kg subcutaneously) 30 minutes before surgeries. Mice were then deeply anesthetized with isoflurane (4% for induction, 1.0–2.5% for maintenance). A 3.5 mm craniotomy was performed above the right hemispheric cerebellum, exposing lobules Crus I and Simplex (6.5 mm anterior to bregma, 2.5 mm lateral to the midline). A window composed of a glass coverslip (Multichannel Systems, CS-3.5R Small round cover glass, #1 thickness) glued (Norland Optical Adhesive 71) to a stainless steel ring (Ziggy’s Tubes and Wires; 316 stainless-steel hypodermic tubing, 9R gauge; OD 0.1470–0.1490″, ID 0.1150–0.1200″; length 0.0197″; cut and deburred) was carefully placed, prefixed with Vetbond (World Precision Instruments), and then glued to the skull with superglue. A one-piece headplate (UCL or Pasteur 3d print facility) was attached around the craniotomy to the animal’s head with dental cement (Paladur, Dentalzon). Metacam (1 mg/kg) was given post-surgery, and animals were allowed at least 3 days of recovery before the start of experiments. In a subset of animals, viral injections and cranial window implantation were performed during a single surgery.

### Two-photon imaging of head-fixed mice *in vivo*

Mice were head-fixed with custom head bars (UCL) to the permanent head plate and rested in a custom plastic tube (10 cm long, 5 cm inner diameter) to reduce the confounding influence of body movements during experiments. Mice were habituated to the set-up and head fixation in at least 3 sessions of at least 15 minutes each before the start of imaging experiments. Air puffs were produced by a Picospritzer III (Parker Hannifin Corporation) and delivered to the whisker (∼1 cm distance, frontal) via a nozzle (1 mm diameter opening) at 5 psi for 1 s, 500 ms or 100 ms. The timing of behavioural images, calcium imaging time series, and whisker air-puff delivery was controlled by triggering TTL pulses with NeuroMatic (*74*) (IGOR Pro, WaveMetrics). Two-photon calcium imaging was performed using a 920 nm laser light generated by a Mai Tai Sapphire laser (Spectra Physics) and a custom-built microscope (2PLSM, Ultima IV, Bruker Nano Surfaces Division, Middleton, WI). The laser beam was scanned across the region of interest through a 16x water-immersion objective (Nikon, 0.8 NA water), which was immersed in ultrasound gel (Sonigel, Mettler Electronics). Excitation power at the back of the objective did not exceed 60 mW. All fluorescence calcium and glutamate transients were imaged in Prairie View using resonant galvo mode. Images were acquired at frequencies between 30 and 232 Hz. Image sizes were 512 x 512 pixels for 30 Hz acquisition rates, and the number of resonant galvo lines was reduced to increase the acquisition rate. A typical experiment consisted of 10 or 30 trials, each trial lasting 15 seconds with stimulation given at 7 seconds (1 s, 0.5 s or 0.1 s whisker air puff). For PF experiments, data were collected within the molecular layer at locations where we observed a high density of labeled PFs. A subset of experiments was performed on a Bruker Ultima 2Pplus using a Vision II laser (Coherent), and a 20x water-immersion objective (Olympus, 1.0 NA), and the timing was controlled using the Prairie View software (Bruker Nano Surfaces Division, Middleton, WI).

#### Cerebellar slice preparation

Acute coronal slices (200 µm thickness) of cerebellum were prepared from adult Gabra6-Cre mice of either sex, aged between P51-P97, for calcium imaging, and from adult CB6F1 mice of either sex, aged between P30 and P90, for synaptic recordings. Following transcardial perfusion with an ice-cold solution containing (in mM): 2.5 KCl, 0.5 CaCl2, 4 MgCl2, 1.25 NaH2PO4, 24 NaHCO3, 25 glucose, 230 sucrose, and 0.5 ascorbic acid, the brains were removed and placed in the same solution. Slices were cut from the dissected cerebellum using a vibratome (Leica VT1200S), and incubated at room temperature for 30 min in a solution containing (in mM): 85 NaCl, 2.5 KCl, 0.5 CaCl2, 4 MgCl2, 1.25 NaH2PO4, 24 NaHCO3, 25 glucose, 75 sucrose, and 0.5 ascorbic acid. Slices were then transferred to an external recording solution containing (in mM): 125 NaCl, 2.5 KCl, 1.5 CaCl2, 1.5 MgCl2, 1.25 NaH2PO4, 24 NaHCO3, 25 glucose, and 0.5 ascorbic acid, and maintained at room temperature for up to 6 hr. All solutions were bubbled with 95% O2 and 5% CO2.

#### Two-photon imaging *in vitro*

The brain slices containing GCaMP8f-expressing GC somata and MF terminals were identified with a 4x objective lens (Olympus UplanFI 4x, 0.13 NA) using very brief illumination with 470 nm light to excite GCaMP8f fluorescence. GCs were identified using infrared Dodt-gradient contrast and a QlClick digital CCD camera (QImaging, Surrey, BC, Canada) mounted on an Ultima multiphoton microscopy system (Bruker Nano Surfaces Division, Middleton, WI, USA) that was mounted on an Olympus BX61W1 microscope, equipped with a water-immersion objective (Olympus 60x, 1.1 NA). Two-photon excitation was performed with a Ti-sapphire laser (Spectraphysics). To visualize GCs expressing GCaMP8f, two-photon excitation was performed at 920 nm. Infrared Dodt-gradient contrast was used to position the stimulation pipette in the white matter, targeting MFs to indirectly activate GC bodies. Linescan imaging of GC somata and MF terminals was performed by scanning through the cell membrane marked by freehand linescan mode (Prairie View). Total laser illumination per sweep lasted 2000 ms. Fluorescence was detected using both proximal epifluorescence and a substage photomultiplier tube gallium arsenide phosphide (H7422PA-40 SEL, Hamamatsu).

#### Synaptic recordings

Whole-cell patch-clamp recordings of GCs were performed using a Multiclamp 700B amplifier (Axon Instruments) and according to the methodology described in Chabrol et al. (*31*). Thick-walled, fire-polished glass patch electrodes (1.5 mm OD, 0.75 mm ID, Sutter Instruments; 5–8 MΩ tip resistance) were filled with a K^+^-based internal solution containing (in mM): 110 KOH, 110 methanesulfonic acid, 6 NaOH, 4.6 MgCl_2_, 5 EGTA, 1.78 CaCl_2_, 4 Na-ATP, 0.3 Na-GTP (280-285 mOsm, pH 7.3). The Crus I or Simplex region was first identified under a 4x objective lens (Olympus UplanFI 4x, 0.13 NA). GCs were then visualized using infrared Dodt-gradient contrast on a multiphoton microscopy system as described above. Cells were voltage-clamped at -80 mV to record EPSCs elicited by electrical stimulation of single MF inputs. MF transmission was evoked using a second patch pipette filled with the recording solution and randomly placed within the GC layer. Single inputs were isolated with 10 μs voltage pulses (Digitimer Ltd, Letchworth Garden City, UK), and stimulation intensity was set to 5 V above the threshold where consistent synaptic responses could be evoked following each pulse in a 100 Hz train. The AMPA-receptor-mediated component of the synaptic current was isolated using SR 95531 (10 μM) to block GABA_A_Rs, d-AP5 (10 μM), and 7-chlorokynurenic acid (20 μM) to block NMDA receptors (NMDARs), and strychnine (0.3 μM) to block glycine receptors. Membrane capacitance and series resistance were estimated online by manual cancellation of the transient current responses to a 5 mV step depolarization. Recordings were performed at 35-37°C, low-pass filtered at 10 kHz, digitized at 100 kHz using an analog-to-digital converter (model NI USB 6259, National Instruments, Austin, TX, USA), and acquired using the Clamp module on NeuroMatic (*77*), running on Igor PRO (Wavemetrics, Lake Oswego, OR, USA).

### EPSCs analysis and estimation of synaptic time constants

Electrophysiological data were analyzed using custom scripts in NeuroMatic (Igor Pro 8). Extracellular stimulation artifacts were subtracted using the Art module, with a single-exponential fit applied to the decay, and baseline correction enabled to account for residual EPSC activity. Series resistance and membrane capacitance were corrected using the online estimates, and traces were low-pass filtered at 4 kHz. Input resistance, estimated from the peak of a small capacitive transient preceding each sweep, was used to monitor cell health. Trials failing the NeuroMatic first-pass stability test for input resistance or for the amplitude of the first EPSC in the train (used as a plasticity proxy) were excluded. Average EPSC amplitude was measured within a 100 μs window centered on the mean peak response. CV was calculated as the square root of baseline-subtracted variance divided by the mean amplitude. PPR was computed as EPSC2/EPSC1 from average 100 Hz trains.

Synaptic decay time constants were estimated by fitting single- and double-exponential models to average EPSC decay waveforms recorded during 20, 100, and 300 Hz stimulation trains. Train durations (3,000 ms at 20 Hz; 500 ms at 100 Hz; 250 ms at 300 Hz) were selected to ensure steady-state responses (yielding 61, 51, and 84 stimuli, respectively). EPSC peaks were detected as absolute minima following each pulse. A custom iterative fitting algorithm selected optimal parameters for each model. Fits were evaluated based on: (1) chi-square values; (2) amplitude ratio of the two components (<10-fold difference); and (3) time constant separation (>2-fold difference). If criteria were met, a weighted decay time constant was calculated from the double-exponential fit as 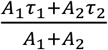. Otherwise, the time constant of a single-exponential model was used. Fits failing to converge—e.g., due to linear EPSC decay—were excluded. Linearity was assessed using the NeuroMatic first-pass stability test and confirmed in GraphPad via slope significance testing.

### Cell-attached recordings and spike analysis

A large-diameter stimulation electrode (>20 μm) was positioned in the white matter to activate MF tracts. Spiking responses were recorded in the cell-attached configuration during 1s trains of 100 Hz stimulation. Pulse width was initially set to 100 μs and adjusted for each cell to optimize activation. Stimulation intensity was increased to maximize the number of MFs recruited, thereby maximizing evoked spike output. When stable access permitted, whole-cell configuration was subsequently established to record the underlying synaptic drive (EPSCs) under identical stimulation conditions. Stimulation artifacts were blanked offline, and spikes were detected using the Spikes module in NeuroMatic to generate peristimulus time histograms (PSTHs). PSTHs were fitted with single- or double-exponential functions to extract spike decay time constants, analogous to what was described for synaptic time constants.

### Data preprocessing

Image registration, Neuropil extraction, movement correction, and cell detection of *in vivo* two-photon imaging data were performed in Suite2p (*78*). For GC and MF recordings, cells were divided into active and inactive groups during the recording session, and only active cells— identified manually based on morphology and activity traces—were further analyzed. Trials were curated for movement artifacts: y-shifts of recording at each timepoint, y-off, as determined by Suite2p, had to be below factor 20. Only cells with a maximum value of the smoothed trace (Savitzky-Golay filter, window length = 9, polynomial order = 2) that exceeded 3 standard deviations of the baseline fluorescence (defined as 1 s before stimulation) during stimulus presentation were considered as responding to the stimulus. Each GC dendritic claw was assigned a unique ID based on morphology and imaging region.

### Imaging data analysis

Data was stored in a MySQL database for further analysis with custom-written Python and MATLAB scripts. Delta F/F_0_ was calculated using a baseline fluorescence period (1 s) immediately prior to stimulation. Calcium traces, 5 seconds long from the start of the stimulus, were fitted by a multi-exponential function with time delay t_0_ one rising *(τ*_*R*_) and two decaying components (*τ*_*D*1_, *τ*_*D*2_) with weights A_1_ and A_2_, respectively (IGOR Pro, WaveMetrics):

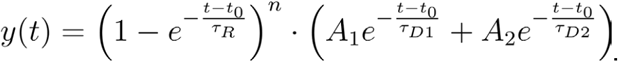

Time to peak, rise time (to 90% of the peak), decay time (time from peak to 50% of maximum), onset time (to 10% above baseline), and FWHM (time from 50% rise to 50% decay) were determined for each cell from their multiexponential fit.

Temporal sparsity was defined as in (*48*):

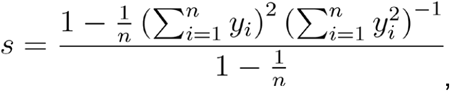

where n is the number of recording time steps.

For *ex vivo* calcium imaging, initial image processing and registration were automated using a custom Python-based pipeline. Individual regions of interest (ROIs) corresponding to GC somata were subsequently extracted from the linescan data using a custom-built viewer in RStudio. During data acquisition, freehand scanning lines were positioned across groups of 5–10 labeled cells per sweep to capture simultaneous activity patterns. To differentiate calcium transients from baseline noise, an automated inclusion criterion was implemented where a cell was considered responsive only if its maximum deflection during the stimulus exceeded 3 standard deviations of the baseline fluorescence. From these traces, we determined the FWHM as described above.

To characterize the specific calcium dynamics (kernels) of individual GC somas, MF terminals, and PF boutons, action potential-evoked transients were recorded in response to 1, 2, and 10 pulses at 100 Hz *ex vivo*. Analysis of these dynamics was performed using Igor Pro and NeuroMatic, where twenty trials were collected for each stimulation condition alongside non-stimulated baseline traces used for subtraction and division of the stimulus trials to generate bleach-corrected ΔF/F_0_ profiles. Kinetic parameters, including peak amplitude, rise time, and 50% decay time, were directly determined from these corrected traces to compare the inherent calcium signaling properties across neuronal compartments.

### Behavioural analysis

Behavioural movies were acquired using two Basler Cameras (ace acA1440-220um USB3 Monochrome, 1440 px x 1080 px resolution), one was filming the mouse head and whiskers at 250 Hz with 500 µs exposure time, the second was filming body movements at 100 Hz. A Python-based custom-written camera driver was used to select optimized image parameters and record single-frame timestamps with microsecond precision. Illumination was provided by an infrared lamp (CMVision IR30 WideAngle IR Illuminator) at 850 nm. ROIs were defined from the video frame to reach the desired recording frequency. Each frame had a unique ID and timestamps up to microsecond precision were automatically recorded. In a subset of experiments, whisker movements were captured at 150 Hz with 500 µs exposure time using a FLIR Blackfly camera (Edmund Optics). To extract whisker kinematics, we used a custom-written Python-based whisker detection and tracking software. The angles of deflection of the left and right whisker arrays were calculated for each image frame within a manually selected radial region around the centre of the whisker pad. Trial-averaged whisker responses to air puff were fitted as for the Ca^2+^ traces using the multi-exponential function described above.

### Histology

Following completion of the behavioral experiments, mice were deeply anesthetized with a mixture of Ketamine (Imalgen, 100 mg/kg, Merial) and Xylazine (Rompun, 10 mg/kg, Bayer) via intraperitoneal injection, and transcardially perfused with PBS, followed by 4% paraformaldehyde (PFA) in PBS. Brains were removed and post-fixed overnight in 4% PFA, then transferred to PBS. Following post-fixation, 50 µm whole-brain sagittal sections were made using a vibratome (Leica Biosystems, V1000S). The sections were then mounted in DAPI (1:5000 dilution) Fluoromount (F4680; Sigma-Aldrich), and the slides were sealed at the margins 24h later with nail varnish and stored at 4° Celsius. Fluorescence images of whole brain slices were captured using a laser-scanning confocal microscope with a 488 nm laser for GFP excitation and a 555 nm laser for tdTomato excitation at 20X magnification of 500 μm^2^ tiles (Axio Scan.Z1, Inverted LSM 700, Zeiss; from the Histology Platform at Institut Pasteur). The images of the brain slices were then matched to the closest brain cartography from the Allen Brain Atlas.

### Statistical analysis

Data plots were generated, and statistical comparisons were performed using the appropriate test in GraphPad Prism version 10.4.1 for Windows (GraphPad Software, Boston, Massachusetts, USA, www.graphpad.com). *P*-values < 0.05 were considered to indicate significance. In summary plots, data are represented as median with interquartile ranges, and box plot whiskers show the full range of values. In figure legends, values indicate median ± standard deviation, except where otherwise noted.

#### Cerebellar cortex model and simulations

Simulations were carried out using a modified version of the cerebellar cortex model developed in (*53*), to which we refer the reader for a detailed description. Below, we give a brief overview and highlight the differences between the original model and the one used here.

#### MF-GC synapses

Model synapses were identical to Barri et al. (*53*), except for the fast refilling time constant (see below). The average synaptic current transmitted per unit time by a single MF to a single GC is:

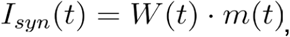

where W(t) is the time-dependent MF-GC synaptic weight and m(t) a continuous variable denoting the instantaneous MF firing rate. W(t) depends on pre-synaptic mechanisms, determining the average number of released vesicles in response to m(t), and a term for post-synaptic receptor desensitization (**Fig. 4B**). Each MF-GC synapse featured two vesicle pools, which differ in their refilling rates and which we refer to as ‘slow’ and ‘fast’, respectively. Thus:

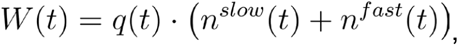

where n^slow^ and n^fast^ are the average number of vesicles released by the two pools, respectively, and q(t) is a post-synaptic factor.

The slow pool was composed of a comparatively small number of docked vesicles, N^slow^, with a high vesicle release probability p^slow,^ and a low rate of recovery from vesicle depletion 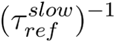 . In addition, the slow pool’s recovery from depletion was augmented by a process of activity-dependent fast vesicle refilling, captured by the probability p_ref_. The fast pool had a comparatively large number of docked vesicles, N^fast^, with a low release probability p^fast^ and a high rate of recovery from depletion 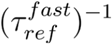. We thus have: 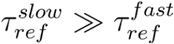, *N*^*slow*^ < *N*^*fast*^ and *p*^*slow*^ > *p*^*fast*^ .

For a single synapse, the fraction of neurotransmitter available at the slow and fast pools, x^slow^ and x^fast^ is given by:

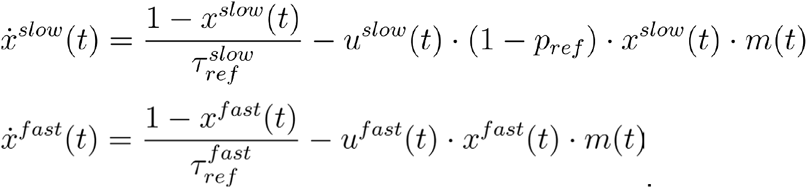

The variables u^slow^(t) and u^fast^(t) denote the pools’ respective release probabilities, which are time-dependent and modified by short-term facilitation:

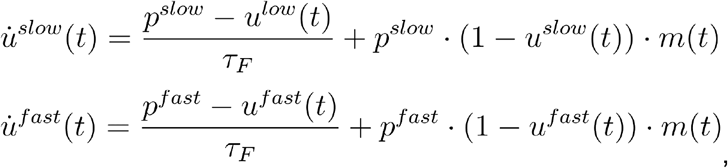

with *τ*_*F*_ denoting the facilitation time-constant. With this, the average number of vesicles released by the two pools can be written as:

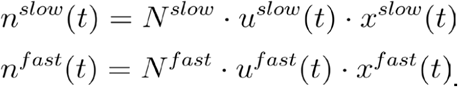

Finally, post-synaptic receptor desensitization depends on the number of released vesicles according to:

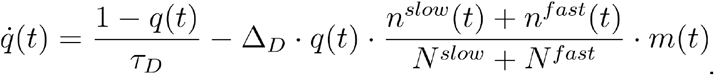

In the absence of STP (**Fig. 4F**), we set *x*^*slow*^ *(t)* = *x*^*fast*^ *(t)* = *q(t)* = 1, *u*^*slow*^ *(t)* = *p*^*slow*^ and *u*^*fast*^ *(t)* = *p*^*fast*^ for all t.

#### Synaptic parameters

As in Barri et al. (*53*), synaptic parameters were constrained to reproduce the average behaviour of the five MF-GC synapse types that were experimentally determined (*31*). All synaptic parameters, except 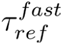, were unchanged from the original model. To obtain region-specific models, we set the fast pool of each synapse to have a 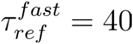 ms for Crus I-like simulations, and 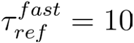 ms for Simplex-like simulations. This resulted in Simplex-like synapses having a higher PPR, longer and higher steady-state amplitudes than Crus I-like synapses, without affecting the initial mean EPCS, in qualitative agreement with our experimental findings (**fig. S7C**). Note that alternatively lowering/increasing release probabilities to obtain Simplex-like/Crus I-like synapses would yield similar qualitative differences in PPR, *τ*_*syn*_ and steady-state amplitudes. However, this would also lead to different initial mean EPSC, which is inconsistent with the experimental data.

#### Mossy Fibers

In our original model, MFs exhibited constant firing rates, dictated by the group membership of their synapses, that experienced an instantaneous frequency switch at stimulus onset (*53*). In contrast, here we modelled air-puff stimulus-evoked activity as multi-exponential functions, with one rising *(τ*_0_) and two decaying components (*τ*_1_, *τ*_2_) to approximate the unobserved firing rates underlying our measured calcium traces. Furthermore, the experimentally observed differences between pontine and Pr5 MFs were qualitatively captured by using two families of firing-rate time profiles.

The rate of the ith (out of M) MF was modelled as:

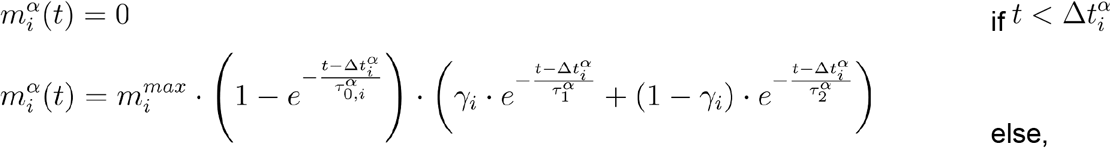

where m^max^ is the peak firing rate, Δ*t* the onset time delay and 0 < *γ* < 1. The index *α* ∈ { pons, Pr5 } indicates pontine or Pr5 origin. MF inputs were associated to the synapse types defined in (*31*) according to the following rule: all synapse types, regardless of their release probability, received pontine MF inputs. Synapses with the highest release probabilities (groups 1 and 2 in (*31*), constituting 22% of all synapses), received Pr5 inputs with probability 0.5, as primary afferents preferentially form high p synapses (*31*).

In keeping with the high firing rates reported in response to whisker stimulation (*56*), peak rates m^max^ were drawn individually for each MF and irrespective of synapse group, and for both pons and Pr5, according to:

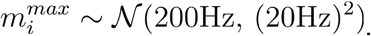

MF onset delays were drawn from exponential distributions

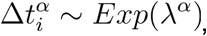

with rate parameters λ^pons^ = 200 ms for pontine MFs and λ^*Pr*5^ = 10ms for Pr5 MFs.

Decay time-constants were set to *τ*_1_ = 100ms, 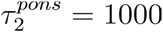 ms and 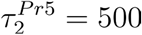 ms to capture the longer duration of pontine MF activation profiles. To account for the temporal variability between MFs, rise time-constants were assigned to be uniformly spaced across M MFs according to 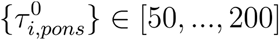 ms and 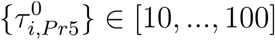 ms. Similarly, we set {*γ*_*i*_} ∈ [0.3, …, 0.9]. To capture the experimental observation that that time to peak was positively correlated with the temporal duration of MF activity (FWHM; **fig. S2**), we introduced rank correlations of 1 between 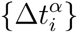 and 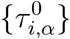 as well as 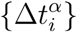 and {*γ*_*i*_}.

#### Cerebellar cortical circuit model

Our model consisted of firing rate units corresponding to M=100 MFs, N=3000 GCs, a single molecular layer interneuron (MLI) and a single PC that linearly summed excitatory inputs from GCs and MLI inhibition. Each GC received four MF synapses randomly selected according to their relative abundance (*31*). The ith GC input h_i_(t) and GC firing rate g_i_(t) were determined by:

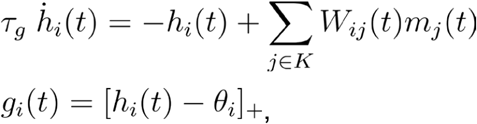

where [⋅]_+_ the linear rectifier, *τ*_*g*_ = 10ms, and K is a set of four MF indices, randomly drawn as explained in (*53*). GC thresholds *θ*_*i*_ were adjusted such that each GC was active for a fraction f_c_=0.5 of all time steps during the interval from stimulus onset to T_post_=1.4 s post-onset. In the absence of STP (**Fig. 4F**), GC thresholds were set identically to those obtained in the presence of STP.

To compute the PC firing rate, each GC firing-rate profile was independently normalized to a peak value of one. MLI activity was assumed to represent the average rate of the GC population, yielding the PC firing rate:

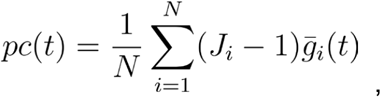

where 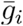 are the normalized GC firing rates and J_i_ are the non-negative GC-PC synaptic weights. Normalizing GC firing rates further simplifies our original model.

Model equations were numerically integrated using the Euler method with step size 0.5 ms.

#### Target signals

Target signals used for PC training were either a realistic PC firing rate profile from (*57*) (**Fig. 4H**) or firing profiles modeled by a gamma distribution according to:

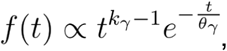

with normalization chosen such that max(f)=1. In **Fig. 4I and 4J** we adjusted *k*_*γ*_ and *θ* _*γ*_ such that the averages of the resulting Gamma shapes corresponded to the target delay and its SDs to 0.01 s and 0.15 s, respectively.

#### Learning rule

Reconstruction of the target signal f(t) was achieved by adjusting the weights J_i_ via a supervised learning rule. The least squares loss to be minimised by the learning procedure is:

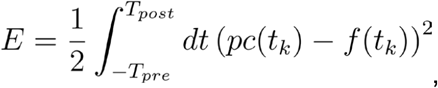

where T_pre_= 0.1 s and T_post_= 1.4 s. Learning via gradient descent leads to a stepwise weight update according to:

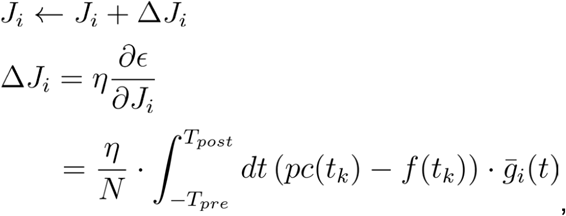

where *η* = 0.0025 is a learning rate. To increase biological realism, we modified the above by relaying the error information through a non-negative and bounded climbing fibre rate:

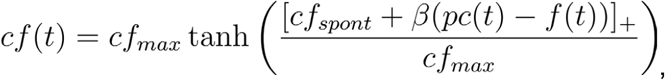

leading to the modified weight update:

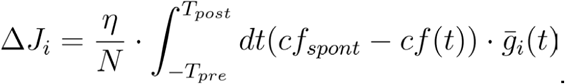

Here, *cf*_*spont*_ = 1 Hz is the spontaneous climbing fibre rate. Relative to the original implementation, we introduced a maximum climbing fibre rate of *cf*_*max*_ = 20 Hz and changed the proportionality factor *β* from 0.5 to 200 to account for the different target signals used here.

As in (*53*), the full gradient update rule (**fig. S9A-C**) and the biologically inspired update rule (**Fig. 4** and **fig. 9D-H**) were modified by a Nesterov acceleration scheme to increase learning speed. For weight learning, we subsampled the simulated and normalized GC rates by a factor of 10 and carried out 4000 update steps. The initial weight distribution was set to J_i_=0 for all i. To calculate the reconstruction error, we computed the final PC output by adding zero-mean white noise with a standard deviation of 0.01 to the normalized GC firing rates.

**Fig. S1.**
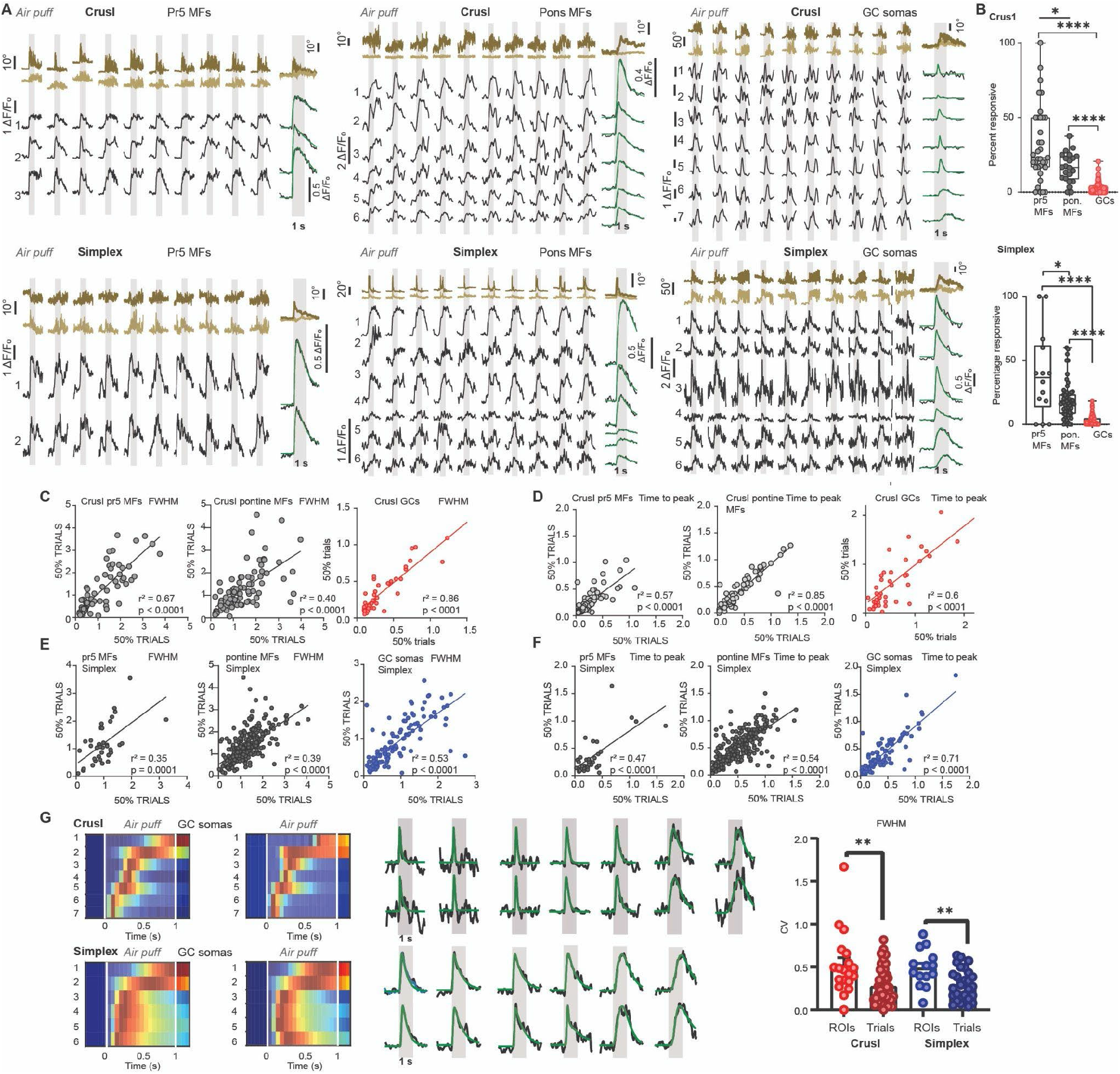
Region-specific response robustness of Pr5 MF, pontine MF, and GC populations. **(A)** Calcium responses in Pr5 mossy fibres (MFs), pontine MFs and granule cell (GC) somata in Crus I and Simplex during ten 1-s whisker stimulation trials. **(B)** Fraction of cells responding to whisker air puff relative to all segmented cells recorded during each experiment. In Crus I, 32 ± 4% of Pr5 mossy fibres (MFs; *n* = 35 experiments, *N* = 4 mice), 16 ± 2% of pontine MFs *(n* = 22, *N* = 3), and 2 ± 0.3% of granule cell (GC) somata *(n* = 132, *N* = 6) responded to whisker air puff. In Simplex, 39 ± 9% of Pr5 MFs *(n* = 14, *N* = 3), 18 ± 2% of pontine MFs *(n* = 58, *N* = 3), and 3 ± 0.4% of GC somata *(n* = 83, *N* = 7) were responsive. **(C-F)**, Correlation of full width at half maximum (FWHM) and time to peak calculated from randomly selected and averaged 50% of trials within each experiment in Crus I (**C,D**) and Simplex (**E,F**). **(G)** Averaged, normalized calcium responses obtained by randomly averaging 50% of trials, respectively, of one representative experiment. Right, the coefficient of variation (CV) is lower within the same cell across trials than across different cells within the same experiment (Simplex, *P* = 0.007; Crus I, *P* = 0.001). Experiments were included in the analysis in which at least 3 cells showed whisker-air-puff responses. Data are presented as median ± interquartile range in the figures. Data are reported in text as mean ± SEM.

**Fig. S2.**
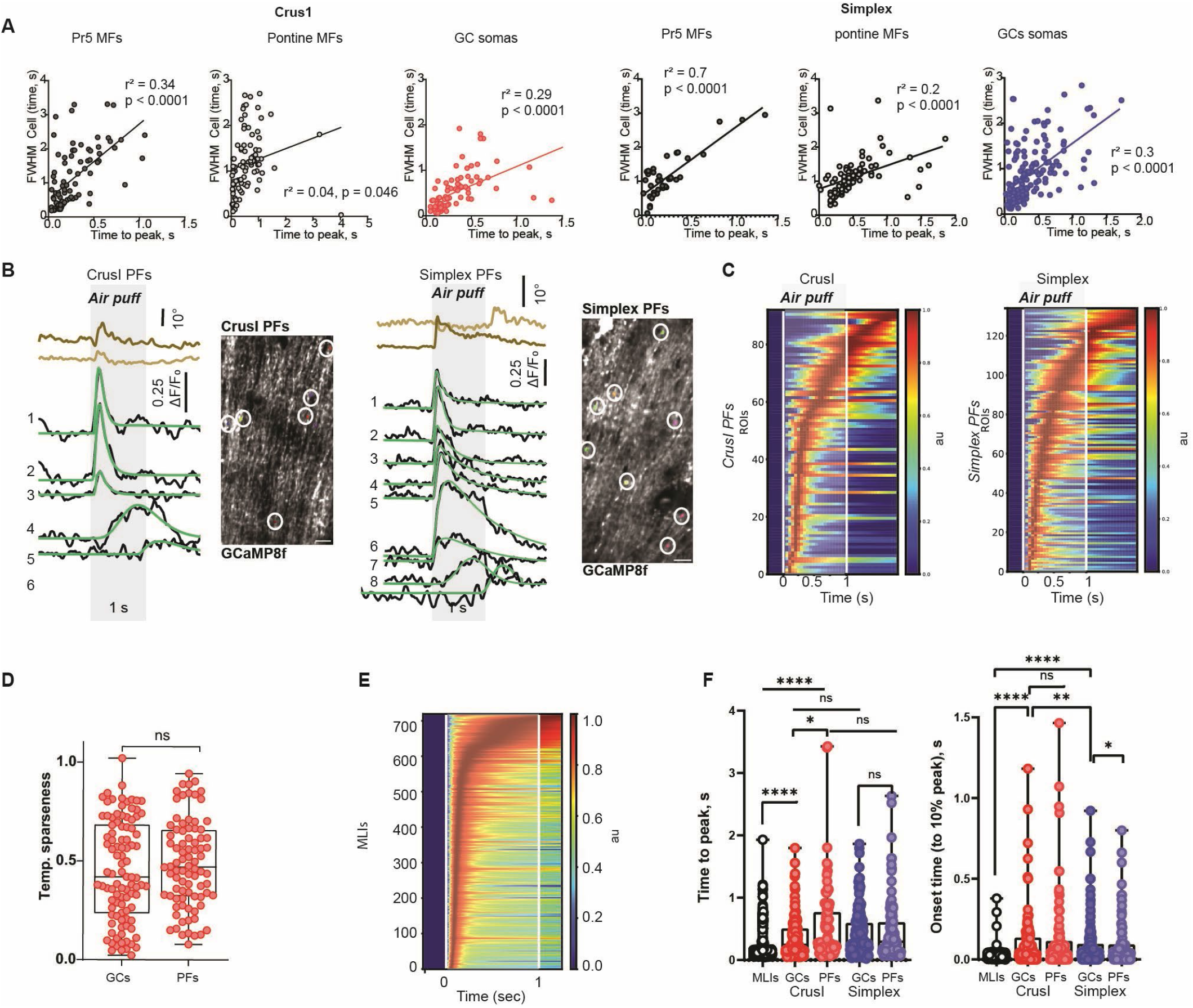
Temporal dynamics of whisker-air-puff-evoked calcium responses across cerebellar inputs, GC somas and PFs, and MLI interneurons. **(A)** FWHM correlated with time to peak in Crus I Pr5 MFs *(n* = 90, *N* = 4), pontine MFs *(n* = 89, *N* = 3), and GCs *(n* = 90, *N* = 5). In Simplex, FWHM also correlated with time to peak in pontine MFs *(n* = 88, *N* = 3), GCs *(n* = 156, *N* = 4), and Pr5 MFs *(n* = 40, *N* = 4). **(B)** Averaged calcium responses of six Crus I granule cell PFs and eight Simplex PFs during ten whisker stimulation trials. **(C)** Raster plots showing PF responses in Crus I *(n* = 92 cells, *N* = 5 mice) and Simplex *(n* = 134, *N* = 6). **(D)** Temporal sparseness did not differ between Crus I GC somata and GC PFs *(P* = 0.316). **e**, Rasterplot of Crus I molecular layer interneuron (MLI) calcium responses to 1-s whisker stimulation *(n* = 720, *N* = 7). **(F)** Time to peak in Crus I MLIs (0.20 ± 0.01 s) was earlier compared with Crus I GC somata (0.36 ± 0.03 s, *P* < 0.0001) and PFs (0.53 ± 0.06 s, *P* < 0.0001). Time to peak did not differ between Crus I and Simplex GCs (0.4 ± 0.03 s, *P* = 0.13) nor between Simplex GCs and PFs (0.45 ± 0.04 s, *P* = 0.92). The time to peak was earlier in Crus I GC somas than in PFs *(P* = 0.015). Onset times were earlier in Crus I MLIs (0.02 ± 0.001 s) compared to Crus I GC somata (0.11 ± 0.01 s, *P* < 0.0001), did not differ between Crus I GC somas and PFs (0.14 ± 0.03 s, *P* = 0.43), were earlier in Simplex GCs (0.08 ± 0.01 s) than Crus I GCs *(P* = 0.01) and did not differ between Simplex PFs (0.09 ± 0.01 s) and Simplex GC somas *(P* = 0.83).

**Fig. S3.**
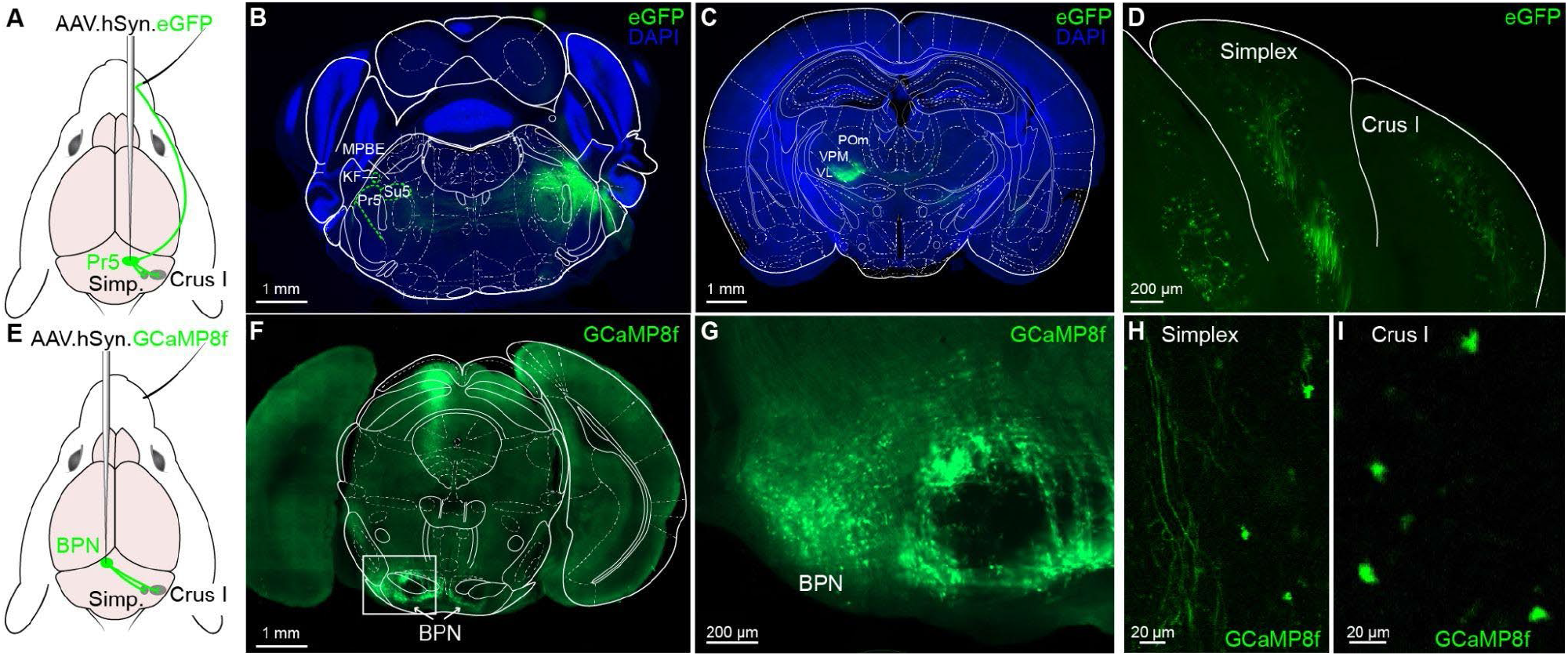
Viral expression in trigeminal nuclei and pons after AAV injections targeting mossy fiber pathways. **(A)** Schematic of AAV.hSyn.eGFP injection into the principal trigeminal nucleus (Pr5) and the resulting projection pattern to cerebellar Simplex and Crus I. **(B)** Coronal section showing eGFP-labeled Pr5 projection fibers (green) in the brainstem. DAPI (blue) outlines anatomical landmarks. Labeled structures include KF, Pr5, and Su5. **(C)** Coronal section showing eGFP expression in thalamic relay nuclei following Pr5 injection. Fluorescence is visible mostly in VPM, the main thalamic target of Pr5. DAPI counterstaining is shown in blue. **(D)** eGFP-labeled mossy fiber terminals in the cerebellar cortex following Pr5 injection. Expression is visible in Simplex and Crus I. White outlines mark lobular boundaries. **(E)** Schematic of AAV.hSyn.GCaMP8f injection into the basilar pontine nuclei (BPN) and the resulting cerebellar projections. **(F)** Coronal section showing GCaMP8f expression in the pons after BPN injection. The white box indicates the region shown in (g). **(G)** Higher-magnification view of the boxed region in (f), showing GCaMP8f-expressing neurons and fibers within the basilar pontine nuclei (BPN). **(H,I)** Mossy fiber terminals in the cerebellar cortex, Simplex and Crus I following BPN injection.

**Fig. S4.**
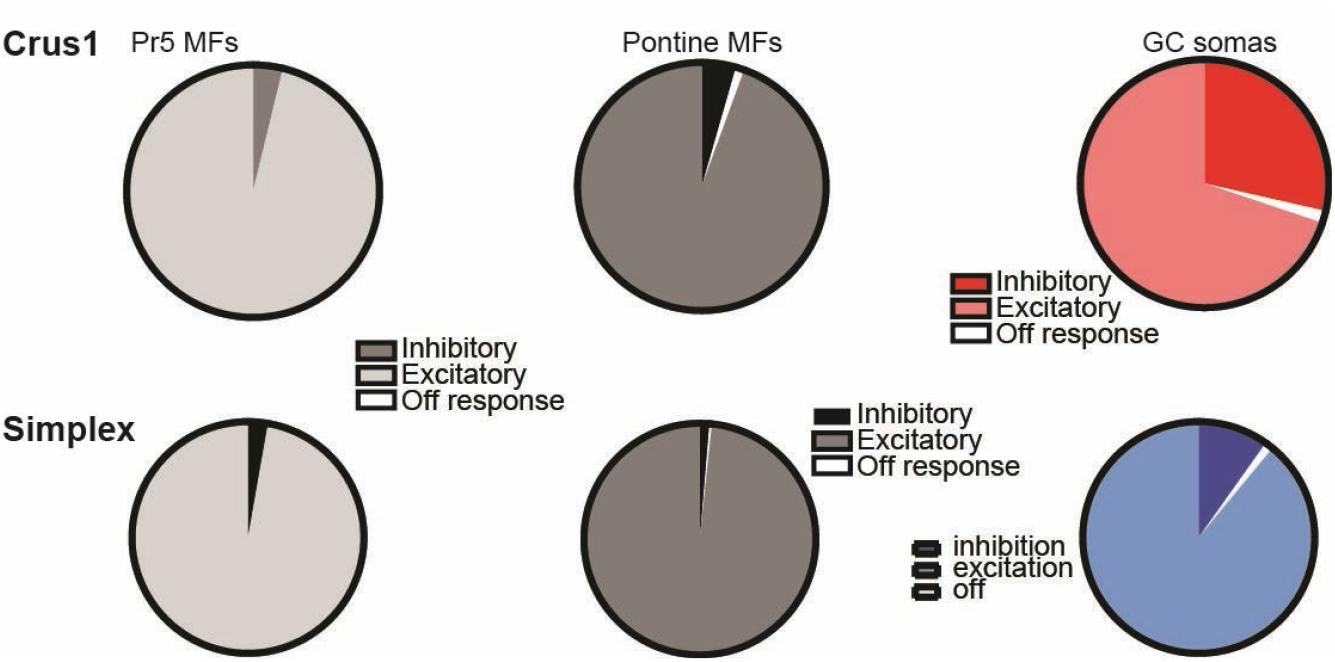
Response polarity distributions. In Crus I, most responses were excitatory in Pr5 MFs (96%), pontine MFs (94%), and GCs (70%), followed by inhibitory responses (Pr5 MFs, 4%; pontine MFs, 5%; GCs, 29%). Off responses were rare (Pr5 MFs, 0%; pontine MFs, 1%; GCs, 2%). In Simplex, 10% of GC responses were inhibitory, and 1% were off responses. In Simplex pontine MFs, 1% of responses were inhibitory, and 0.5% were off responses, whereas in Pr5 MFs, 3% were inhibitory, and no off responses were observed.

**Fig. S5.**
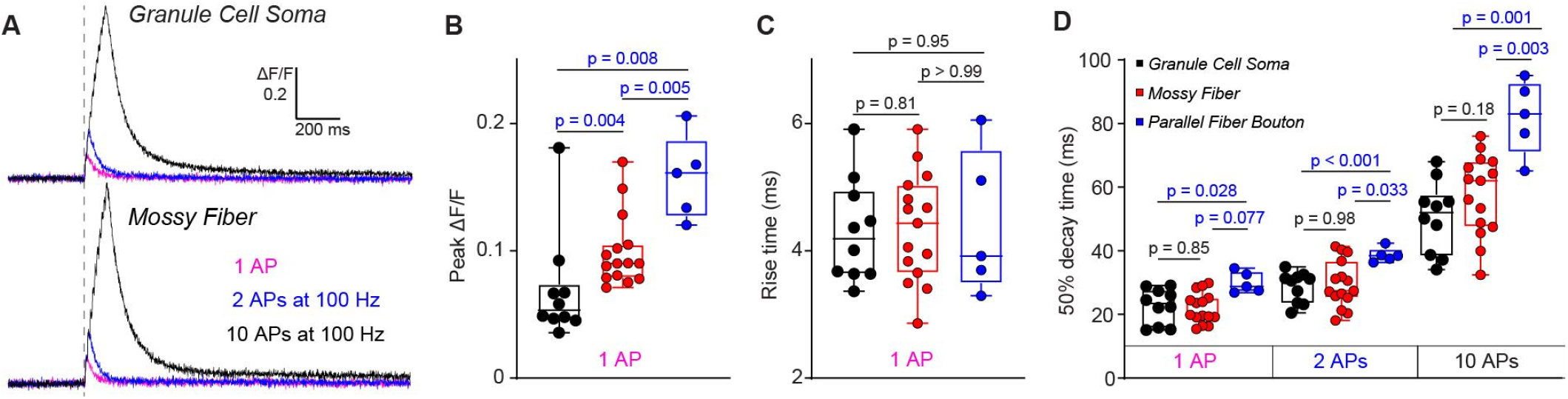
GCaMP8f calcium signals in granule cell somata, mossy fibers, and parallel fiber boutons. **(A)** Example calcium transients recorded from a granule cell soma (top) and a mossy fiber bouton (bottom) in response to 1 action potential (AP; magenta), 2 APs at 100 Hz (blue), and 10 APs at 100 Hz (black). The ΔF/F scale and time scale are indicated. **(B)** Peak ΔF/F values for granule cell somata (median = 0.05, *n* = 10), mossy fibers (0.09, *n* = 15), and parallel fiber boutons (0.1, *n* = 5) in response to 1 AP. Statistical comparisons (Mann–Whitney tests): soma vs mossy fiber *(P* = 0.004); mossy fiber vs bouton *(P* = 0.005); soma vs bouton *(P* = 0.008). (**C)** Rise times for granule cell somata (median = 4.2 ms, *n* = 10), mossy fibers (4.4 ms, *n* = 15), and parallel fiber boutons (3.9 ms, *n* = 5) in response to 1 AP. Statistical comparisons (Mann–Whitney tests): soma vs mossy fiber *(P* = 0.8); mossy fiber vs bouton *(P* > 0.99); soma vs bouton *(P* = 0.95). **(D)** 50% decay times measured for 1 AP, 2 APs at 100 Hz, and 10 APs at 100 Hz. 1 AP: granule cell somata (median = 23.5 ms, *n* = 10), mossy fibers (19.9 ms, *n* = 15), and parallel fiber boutons (28.8 ms, *n* = 5). Statistical comparisons (Mann–Whitney tests): soma vs mossy fiber *(P* = 0.85); mossy fiber vs bouton *(P* = 0.008); soma vs bouton *(P* = 0.03). 2 APs at 100 Hz: somata (30.9 ms, *n* = 10), mossy fibers (27.4 ms, *n* = 15), and boutons (38.5 ms, *n* = 5). Statistical comparisons: soma vs mossy fiber *(P* = 0.98); mossy fiber vs bouton *(P* = 0.03); soma vs bouton *(P* = 0.0007). 10 APs at 100 Hz: somata (52.1 ms, *n* = 10), mossy fibers (62.1 ms, *n* = 15), boutons (83 ms, *n* = 5). Statistical comparisons: soma vs mossy fiber *(P* = 0.18); mossy fiber vs bouton *(P* = 0.003); soma vs bouton *(P* = 0.001).

**Fig. S6.**
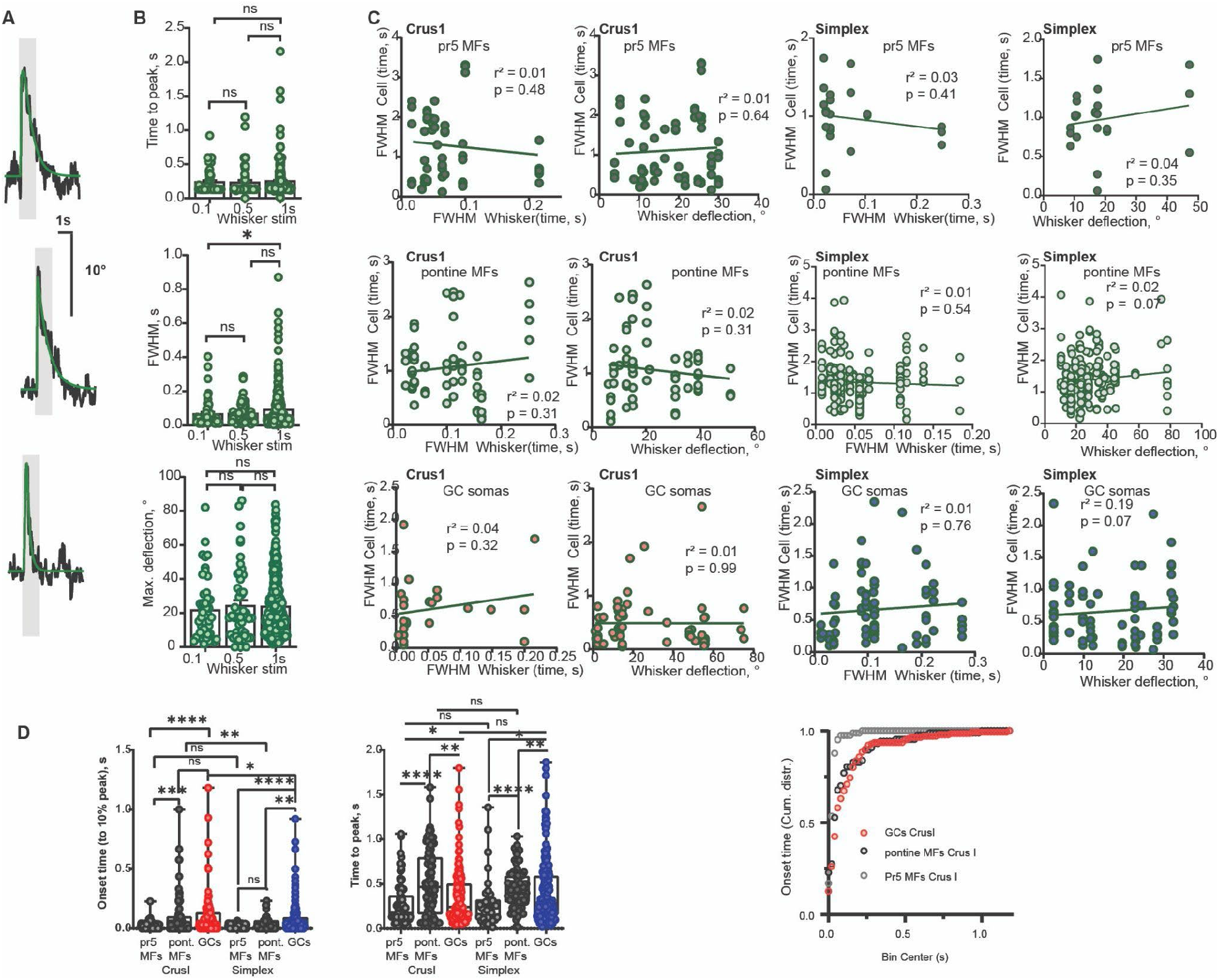
Whisker kinematics and timing of cerebellar MG and GC responses to whisker air-puff stimulation. **(A)** Representative curve fits of contralateral whisking response to 1s whisker air puff stimulation, averaged across trials. **(B)** Time to peak of averaged over trials whisker response of the whisker contralateral to the air puff was not different for 1 s (0.25 ± 0.01 ms, *N* = 29 mice, *n* = 379 experiments) compared to 100 ms (0.24 ± 0.02 s, *N* = 17, *n* = 59, *P = 0*.*6)*, or 500 ms (0.23 ± 0.03 s, *N* = 18, *n* = 54, *P = 0*.*34)* whisker air puff stimulation. FWHM of Whisker response was not different between 0.1 s (0.06 ± 0.01s, *n* = 51, *N* = 17) and 0.5 s (0.07 ± 0.01 s, *n* = 50, *N*= 17, *P = 0*.*15)* or 0.5 and 1 s (0.09 ± 0.01 s, *n* = 340, *N* = 31, *P = 0*.*48)*, but slightly longer for 1 s compared to 0.1 s *(P = 0*.*02)*. Maximal whisker deflection was not different for 1 s whisker air puff stimulation (24.02 ± 0.93°, *N* = 27, *n* = 330) compared to 100 ms (21.88 ± 2.62°, *N* = 15, *n* = 48, *P = 0*.*2)* or 500 ms (24.43 ± 3.06°, *N* = 17, *n* = 50, *P = 0*.*51)*. **(C)** FWHM of whisker response did not correlate with FWHM of CrusI Pr5 MF, pontine MF or GC responses. Maximal deflection of whisker response did not correlate with FWHM of CrusI pr5 MF *(N* = 4 mice, *n* = 76 cells), pontine MF *(N* = 3, *n* = 86) or GC *(N* = 5, *n* = 83) responses. FWHM of whisker response did not correlate with FWHM of Simplex GCs, pontine MF, or Pr5. **(D)** Response onset was delayed in Crus I GCs (0.1 ± 0.014 s) compared with Pr5 MFs (0.03 ± 0.005 s, *P* < 0.0001) but not significantly compared with pontine MFs (0.1 ± 0.02 s, *P* = 0.2). Response onset was later in Simplex GCs (0.08 ± 0.009 s) than in pontine MFs (0.04 ± 0.005 s, *P* = 0.0018) or Pr5 MFs (0.02 ± 0.003 s, *P* < 0.0001). Time to peak was later in Crus I pontine MFs (0.51 ± 0.04 s) than in Pr5 MFs (0.27 ± 0.02 s, *P* = 0.0006) or GCs (0.36 ± 0.03 s, *P* = 0.002). Also in Simplex, time to peak was later in pontine MFs (0.43 ± 0.02 s) than in Pr5 MFs (0.29 ± 0.04 s, *P* < 0.0001) or GCs (0.41 ± 0.03 s, *P* = 0.007). Right, cumulative distribution of Onset time for GCs, pontine MFs, and Pr5 MFs. Note the overlap of GCs and pontine MFs.

**Fig. S7.**
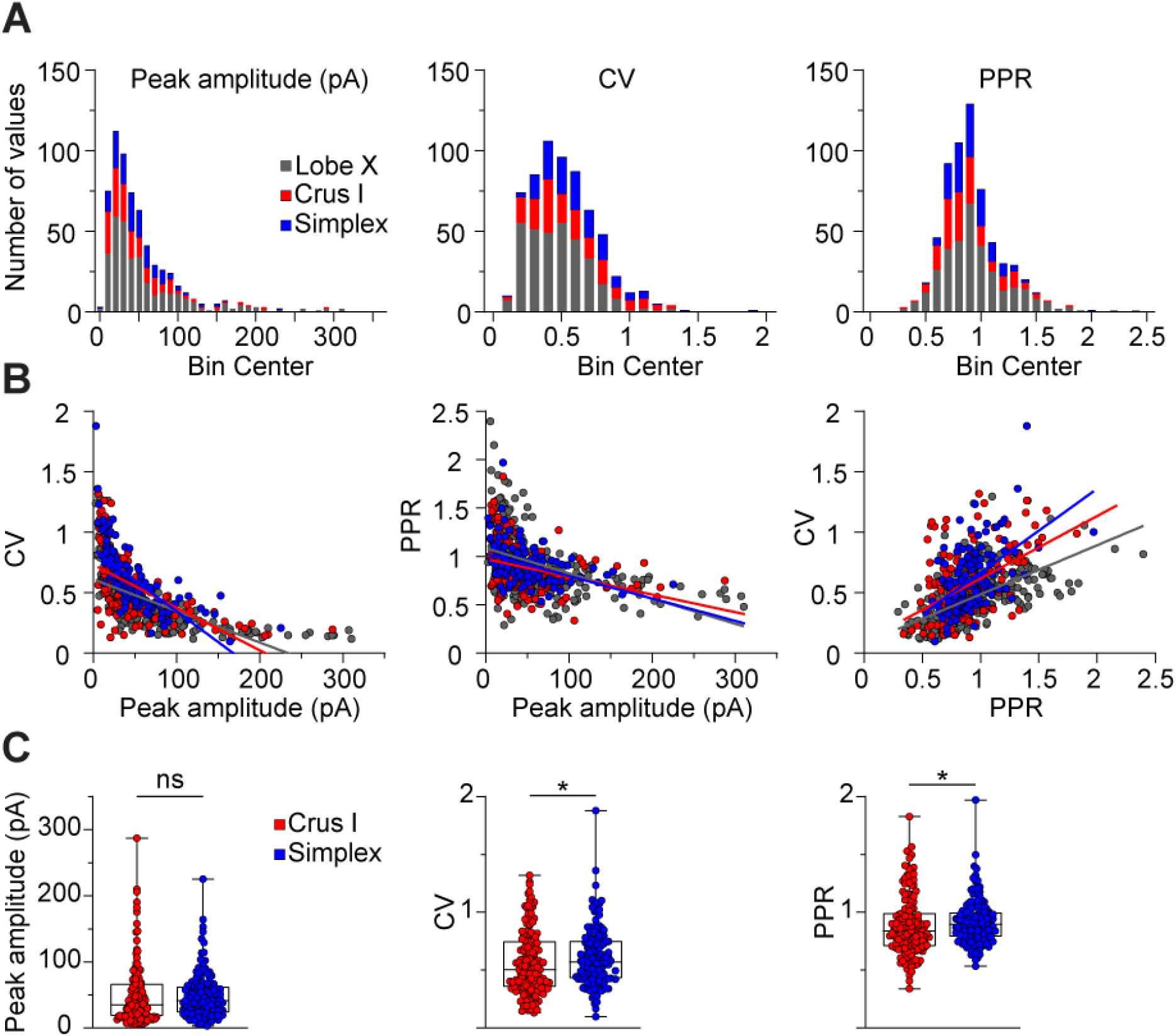
Diverse MF-GC synaptic response metrics in Crus I and Simplex mirror vestibulocerebellar heterogeneity but vary between regions. **(A)** Stacked frequency distributions of EPSC peak amplitude (Lobe X: 37.50 ± 55.60, Crus I: 34.97 ± 45.19, Simplex: 41.30 ± 34.17), coefficient of variation of peak amplitude (CV; Lobe X: 0.45 ± 0.21, Crus I: 0.50 ± 0.27, Simplex: 0.57 ± 0.25), and paired-pulse ratio (PPR; Lobe X: 0.89 ± 0.31, Crus I: 0.84 ± 0.26, Simplex: 0.90 ± 0.20) at 100 Hz stimulation from 326 MF-GC synapses recorded in Lobe X (vestibulocerebellum, see Chabrol et al., 2015), 158 synapses in Crus I and 143 synapses in Simplex using a blind MF stimulation protocol. Distribution shapes are similar across areas, suggesting similar diversity in STP dynamics at MF-GC synapses. **(B)** Correlation plots between EPSC peak amplitude, CV and PPR across the recorded synapses in each region, with corresponding regression lines (Peak amplitude vs CV; Lobe X: *R*^*2*^ = 0.46, Crus I: *R*^*2*^ = 0.34, Simplex: *R*^*2*^ = 0.47; Peak amplitude vs PPR; Lobe X: *R*^*2*^ = 0.22, Crus I: *R*^*2*^ = 0.10, Simplex: *R*^*2*^ = 0.16; PPR vs CV; Lobe X: *R*^*2*^ = 0.34, Crus I: *R*^*2*^ = 0.23, Simplex: *R*^*2*^ = 0.29). Significant correlations were detected across all parameters in each region *(P* < 0.0001 for all pairs, Spearman correlation). (**C**) Quantification of peak amplitude (left; Crus I: 34.97 ± 45.19 pA, Simplex: 41.30 ± 34.17 pA), CV (middle; Crus I: 0.50 ± 0.27, Simplex: 0.57 ± 0.25), and PPR (right; Crus I: 0.84 ± 0.26, Simplex: 0.90 ± 0.20). Simplex synapses show higher CV *(P* = 0.024, Mann-Whitney test) and PPR *(P* = 0.024, Mann-Whitney test) (Crus I: *n* = 158, Simplex: *n* = 143).

**Fig. S8.**
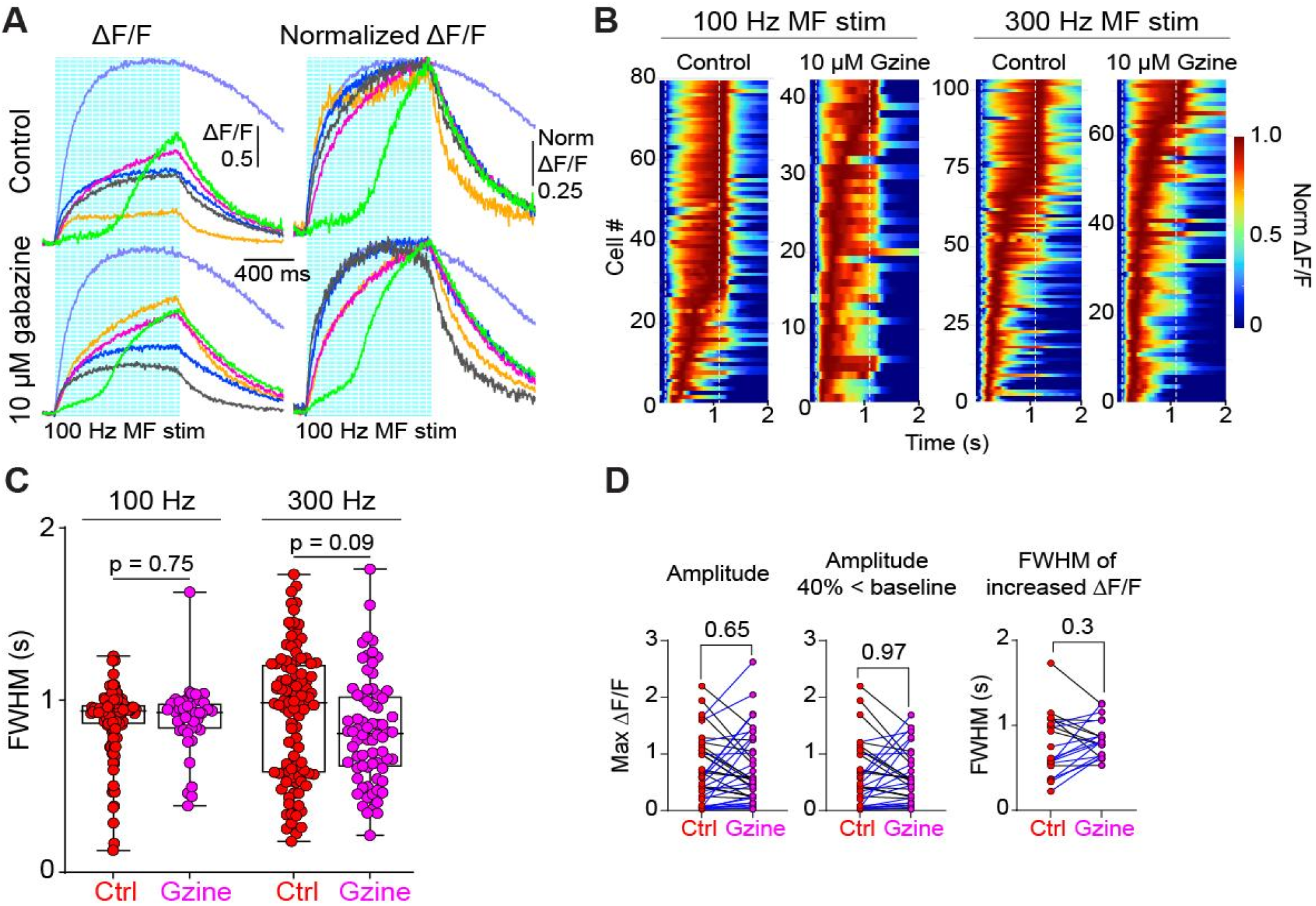
Effect of GABA_A_ receptor blockade on GC calcium responses during MF stimulation. **(A)** Example ΔF/F calcium traces (left) and normalized ΔF/F traces (right) recorded from granule cells during 100 Hz MF stimulation under control conditions (top) and in the presence of 10 μM gabazine (bottom). Individual cells are color-coded. Vertical blue lines indicate the timing of each MF stimulus pulse. **(B)** Raster plots of normalized ΔF/F signals from simultaneously recorded granule cells during 100 Hz (left) and 300 Hz (right) MF stimulation under control conditions and after application of 10 μM gabazine. Each row represents a single cell. Dashed white lines mark the onset and offset of the MF stimulation. Color scale indicates normalized ΔF/F. (**C)** Full width at half maximum (FWHM) of granule cell calcium responses during 100 Hz (left) and 300 Hz (middle) MF stimulation under control conditions and with gabazine. Box plots show the median and interquartile range; each point represents a single cell (100 Hz: control median = 0.94 s, *n* = 117; gabazine median = 0.93 s, *n* = 42. 300 Hz: control median = 0.98 s, *n* = 104; gabazine median = 0.81 s, *n* = 72; Mann–Whitney test *P* = 0.09). **(D)** Summary of granule cell calcium response measurements under control conditions and after application of 10 μM gabazine. The left plot shows the peak ΔF/F amplitude for all paired cells (Wilcoxon matched-pairs signed-rank test, *P* = 0.65; n = 42). The middle plot shows peak ΔF/F amplitude after excluding cells whose baseline fluorescence increased by more than 40% during gabazine application, used as a control to ensure that effects were not driven by baseline shifts (Wilcoxon matched-pairs signed-rank test, *P* = 0.97; n = 35). The right plot shows the FWHM measured only from cells whose ΔF/F increased during gabazine application (Wilcoxon matched-pairs signed-rank test, *P* = 0.33; n = 20).

**Fig. S9.**
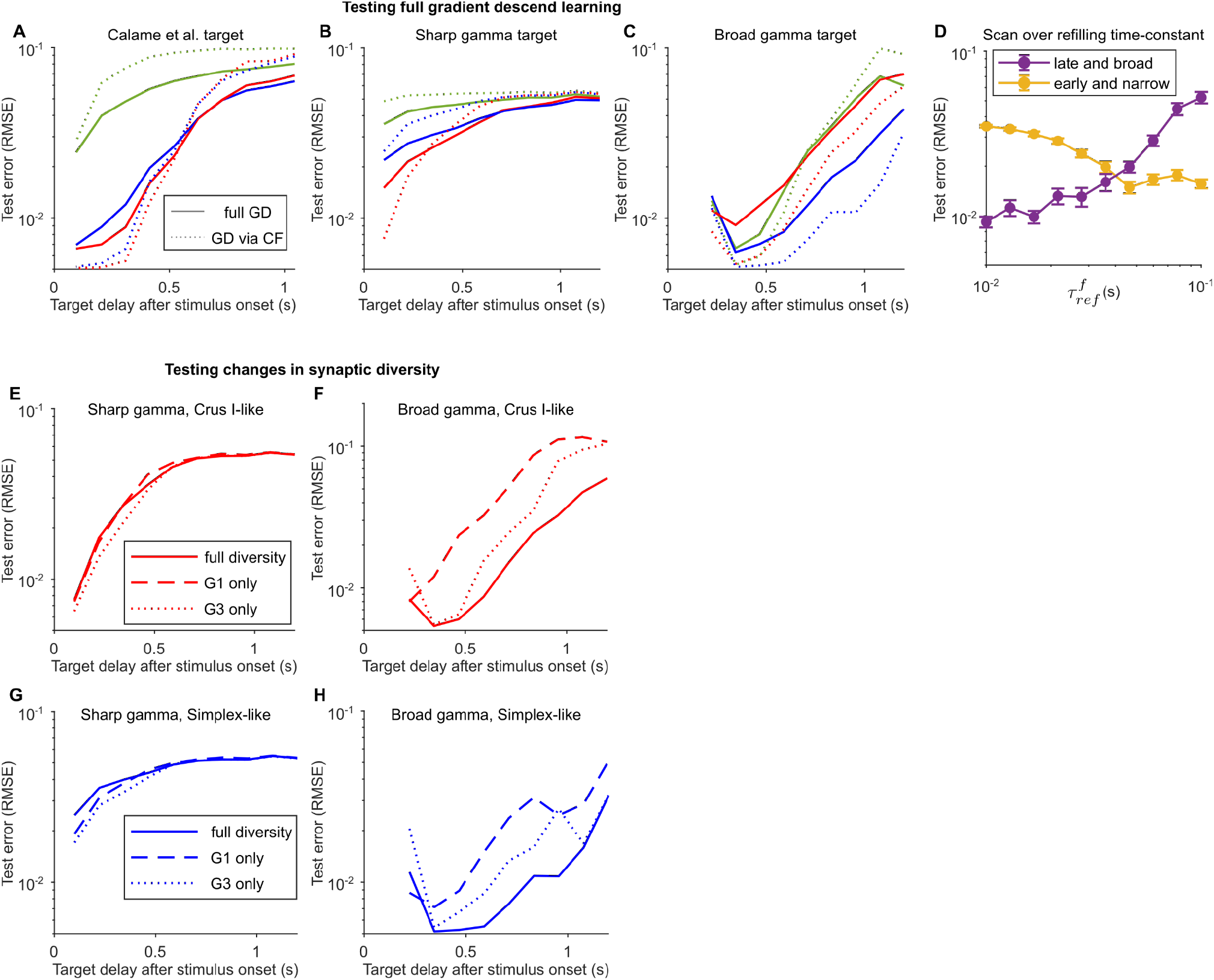
Full vs restricted gradient descent learning, role of synaptic refilling time-constant, and effect of synaptic diversity on reconstruction error. **(A)** Reconstruction error of a sharp temporal signal representing forelimb reach kinematics (Calame et al., 2023) at varying target delays for different GC basis sets with biologically inspired learning rule (dotted lines) and full gradient descent (solid lines); each point represents an average over 30 learning simulations. **(B)** Same as (**A**), but for narrow gamma target signals with a SD of 0.01 s. **(C)** Same as (**B**), but for broad gamma target signals with a SD of 0.15 s. **(D)** Reconstruction error of gamma target signals with a delay of 0.25 s and with a SD of 0.01 s (yellow line) and with a delay of 0.75 s and with a SD of 0.15 s (purple line), for varying fast vesicle refilling time-constants. **(E)** Reconstruction error of gamma target signals with a SD of 0.01 s at varying target delays for Crus I-like models; different lines denote models with varying degrees of synaptic diversity: full diversity (solid line), all synapses assigned to high release probability group 1 (dashed line), and all synapses assigned to low release probability group 3 (dotted line). **(F)** Same as (E) but for broad gamma target signals with a SD of 0.15 s. **(G,H)** Same as (**E,F**) but for the Simplex-like model.

**Fig. S10.**
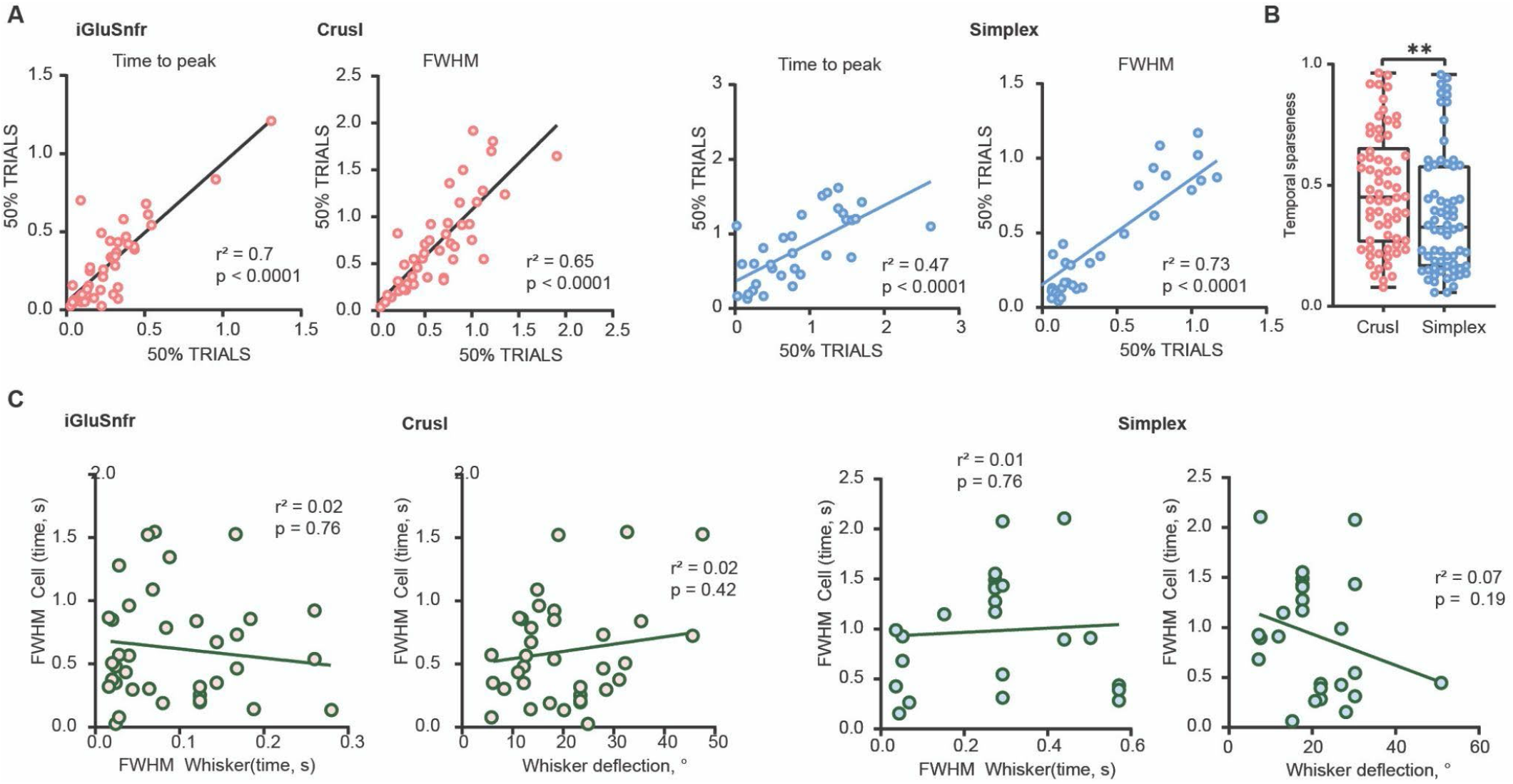
Reliable and region-specific temporal dynamics of MF–GC synaptic responses to whisker airpuff. **(A)** GCs were sparsely labelled with iGluSnFR under the TRE promoter in TCGO mice. Correlations calculated from randomly averaged 50% of trials, respectively, for MF–GC synapses in Crus I and Simplex for time to peak and FWHM. **(B)** MF–GC synaptic responses were more temporally sparse in Crus I (0.48 ± 0.03 s) than in Simplex (0.38 ± 0.03 s, *P* = 0.008). **(C)** No correlation was observed between MF–GC synaptic temporal dynamics and responses to contralateral whisker stimulation in experiments in which the average whisker response could be fitted to a multiexponential function *(N* = 9, *n* = 58; Simplex, *N* = 8, *n* = 40).

## Notes

### Competing Interest Statement

The authors have declared no competing interest.

### Summary of Updates

This manuscript version contains an updated abstract, introduction, and methods.

